# A new subgenome of the *Camelina* genus reveals genome dominance is controlled by chromosomal proximity

**DOI:** 10.1101/2025.08.18.670935

**Authors:** Raju Chaudhary, Kevin C. Koh, Peng Gao, Sampath Perumal, Erin E. Higgins, Kyla Horner, Stephen J. Robinson, Zhengping Wang, Christina Eynck, Venkat Bandi, Andrew G. Sharpe, Isobel A. P. Parkin

## Abstract

*Camelina sativa* is an oilseed of the *Brassicaceae*, whose close relatives vary in ploidy number, providing a novel platform for studying plant genome evolution. The availability of diploid, tetraploid and hexaploid species of *Camelina* allow the evolutionary trajectory and fate of duplicated genes in the neopolyploid *Camelina* species to be elucidated. Here we report an improved assembly of the widely used *C. sativa* reference DH55 and three new genome assemblies of *Camelina microcarpa*; one tetraploid CN119243 (2n = 26), and two hexaploids with divergent chromosome numbers, Type 1 - CN119205 (2n=40) and Type 2 - CN120025 (2n=38). The tetraploid represents the first step in the evolutionary path to form *C. sativa*, while the hexaploids suggest three divergent lineages in the formation of higher ploidy *Camelina* species. The previously uncharacterized fourth subgenome found in *C. microcarpa* Type 2, although showing some homology to the *C. sativa* diploid progenitor genome, *C. neglecta*, showed numerous unique chromosomal rearrangements differentiating it from other subgenomes present in known *Camelina* species. Although this species was recently formed, the second subgenome showed gene expression dominance, which was in contrast to both 2n=40 *Camelina* species where the third subgenome was dominant. The expression dominance in Type 2 *C. microcarpa* contradicted the accepted two-step evolutionary process which led to the generation of related Brassicaceae species. However, the observed genome dominance in all *Camelina* species was negatively correlated with inter-subgenome chromatin interaction frequencies, suggesting that chromosome confirmation and proximity in the nucleus contributes to this mechanism of genome evolution. Despite the differences in genome structure, successful inter-specific hybridization provided evidence of chromosomal exchange between the divergent third sub-genomes of *C. sativa* and *C. microcarpa* Type 2, opening up a novel avenue to new diversity in the established oilseed.

**Key points:**

- An improved genomic understanding of *Camelina* species and identification of distinct subgenome structures and relationships, which will facilitate strategies to increase the genetic diversity in *C. sativa*.
- Subgenome evolution and subgenome dominance in polyploids is associated with chromosomal architecture and proximity in the nucleus.
- Genome assemblies representing all ploidy levels in the *Camelina* genus provide a unique and valuable platform for polyploid research.

## Introduction

*Camelina sativa* (L.) Crantz is a hexaploid species of the Brassicaceae family (Al-Shehbaz et al., 2006). Due to a unique fatty acid composition as well as protein content, *C. sativa* has shown economic potential as an ingredient for the aquaculture, poultry feed and cosmetics industries, as well as being favoured for human consumption (Berti et al., 2016). One factor limiting crop improvement in *C. sativa* has been the low natural genetic diversity within the species (Ghamkhar et al., 2010; Luo et al., 2019; Singh et al., 2015).

In other crops, related or wild species have shown potential as sources of novel variation (Tirnaz et al., 2022); in this context, identification of diverse haplotypes for regions of interest in related *Camelina* species, coupled with suitable genomic resources might be useful in furthering work in *C. sativa* (Brock et al., 2020). Several wild relatives of *C. sativa* have been reported, with some well-defined taxonomically, while others show evidence of cryptic species with comparable morphologies (Martin et al., 2017). *Camelina microcarpa* Andrz. ex DC is one of the closest wild relatives of this crop and was believed to encompass three levels of ploidy; diploid, tetraploid and hexaploid (Brock et al., 2018). For the latter level, two different hexaploid *C. microcarpa* species with distinct combinations of three subgenomes have been now identified (Chaudhary et al., 2020). The first hexaploid *C. microcarpa* appears to share three subgenomes with *C. sativa* (hereafter *C. microcarpa* “Type 1” or CmiT1), whereas the second, *C. microcarpa* (hereafter *C. microcarpa* “Type 2” or CmiT2) shares only two subgenomes with *C. sativa*, while the third subgenome is distinct from any of the three subgenomes of *C. sativa* (Chaudhary *et al*., 2020). A number of whole genome duplication events and chromosomal rearrangements seem to have occurred in the generation of the higher ploidy *Camelina* species. The characterization of the subgenome composition in *Camelina* species will provide a strategy for interspecific hybridization in *C. sativa* and could potentially help in broadening the narrow genetic base of *C. sativa* germplasm (Brock *et al*., 2018; Luo *et al*., 2019; Singh *et al*., 2015).

After whole-genome hybridisation in an allopolyploid the orthologous duplicate genes commonly show evidence of higher gene expression for a particular subgenome (genome dominance) and gene loss (fractionation), which is generally more prevalent in the less dominant subgenome. Such observations led to the two-step-hypothesis in the formation of Brassica paleo-hexaploids (Lyons et al., 2008), whereby the addition of each subgenome led to inferred dominance over the resident genome, generating a hierarchy of gene loss from the Least Fractionated (LF) to the Most Fractionated (MF) subgenomes (Tang et al., 2012). Although gene fractionation is relatively low in *C. sativa* (Kagale et al., 2014) likely due to its recent emergence as a polyploid, there are marked differences in gene expression among the orthologs and the third subgenome was generally found to be dominant (Chaudhary *et al*., 2020; Kagale et al., 2016). The exact mechanism controlling subgenome dominance is not known, although chromosomal rearrangements such as homoeologous exchanges or differences in chromosome architecture such as the level of transposons have both been associated with the observed effect on gene expression (Bird et al., 2019; Edger et al., 2019; Woodhouse et al., 2014). Likewise, epigenetic and structural changes after genome merger could also play a role in shaping gene expression in the polyploids (Cheng et al., 2016; Hollister and Gaut, 2009). However, there are species which have either undergone limited or unbiased fractionation after genome duplication such as *Cucurbita* and soybean, respectively (Sun et al., 2017; Zhao et al., 2017). For *C. microcarpa* T2, the lack of the third dominant subgenome of *C. sativa* presents an interesting candidate to explore the nature of subgenome dominance. Further, the availability of genome assemblies for diploid *C. neglecta* (Chaudhary et al., 2023b) and diploid *C. hispida* (Boiss.) Hedge (Martin et al., 2022), both posited as representative progenitors for *C. sativa,* provides an opportunity to explore the evolution of genome structure in higher ploidy Camelina species.

Here, we sequenced the genome of tetraploid *C. microcarpa* (CN119243) (2n = 26), hexaploid *C. microcarpa* T1 (CN119205) (2n = 40), hexaploid *C. microcarpa* T2 (accession CN120025) (2n=38) and hexaploid *C. sativa* (accession DH55); which revealed the subgenome structure of a previously uncharacterised fourth subgenome for the *Camelina* genus. The relationship among subgenomes was explored in terms of age of divergence, syntenic genomic block rearrangements and chromatin interactions. Gene expression dominance was observed in all three hexaploid *Camelina* species, where the non-dominant genomes showed higher numbers of chromosomal interactions, suggesting reduced proximity of the dominant genome allowed independent and hence greater gene expression.

## Results

### De novo genome assembly, repeat analysis, and gene annotation of *Camelina* genomes

The sequencing of the *C. microcarpa* and *C. sativa* genomes used both Pacific BioSciences (PacBio) and Oxford Nanopore Technology (ONT) long read sequencing techniques. *Camelina microcarpa* CN119243 (Cmi4X) is a tetraploid, whereas the other three genotypes CN120025 (CmiT2), CN119205 (CmiT1) and DH55 (CsaDH55)) are hexaploid in nature. Initial genome assembly was performed with 12-106× coverage of ONT, 14-36× PacBio HiFi, and 32× of PacBio CCS reads (**Supplementary Table 1),** which resulted in base assemblies with contig N50 lengths ranging from 3.4 Mb to 19.82 Mb. Further, the base assemblies were scaffolded using chromosome conformation capture sequencing data (HiC) to generate contiguous assemblies with scaffold N50 lengths of 26.09 Mb, 29.88 Mb, 28.42 Mb, and 32.12 Mb for Cmi4X, CmiT2, CmiT1, and CsaDH55, respectively (**Supplementary Fig. 1**). The genomes comprised either 13, 19, or 20 pseudomolecules, respectively as expected based on the known chromosome numbers. The chromosomes were assigned to subgenomes (SG) and numbered according to the sequence similarity shown after alignment with the *C. neglecta* and *C. sativa* genomes as described in Chaudhary *et al*. (2020); such that SG1 (chromosome 1 - 6) is similar to both *C. neglecta* and the first subgenome of *C. sativa*, SG2 (chromosome 7 - 13) is similar to the second subgenome of *C. sativa*, and SG3 (chromosome 14 - 19/20) is the remainder of the genome (**Fig. 1**). The quality of the genomes were confirmed by various methods, including BUSCO analysis (Simão et al., 2015) which identified between 99.2% and 99.5% of the Embryophyta universal single-copy orthologues in each of the genomes (**Table 1**). Further, Qualimap v.2.2.2 (Okonechnikov et al., 2016) analysis indicated more than 99.42% of available Illumina reads were mapped to the genome with a general error rate of 0.0027% (**Supplementary Table 2**). The assembled genomes comprised more than 91.04% of the estimated genome size using kmer analysis (Marçais and Kingsford, 2011) (**Supplementary Fig. 2**). The assembled genomes contain more than 93.76% of the genomic sequences in pseudomolecules or chromosomes for Cmi4X, 99.34% for CmiT2, 96.63% for CmiT1, and 96.38% for DH55. The representation of full length long terminal repeat (LTR) elements was high for each of the genomes as measured by the LTR assembly index (LAI) (Ou et al., 2018), ranging between 18.27 and 22.49, suggesting high quality assembly of the repetitive fraction of the genome.

**Figure 1.**
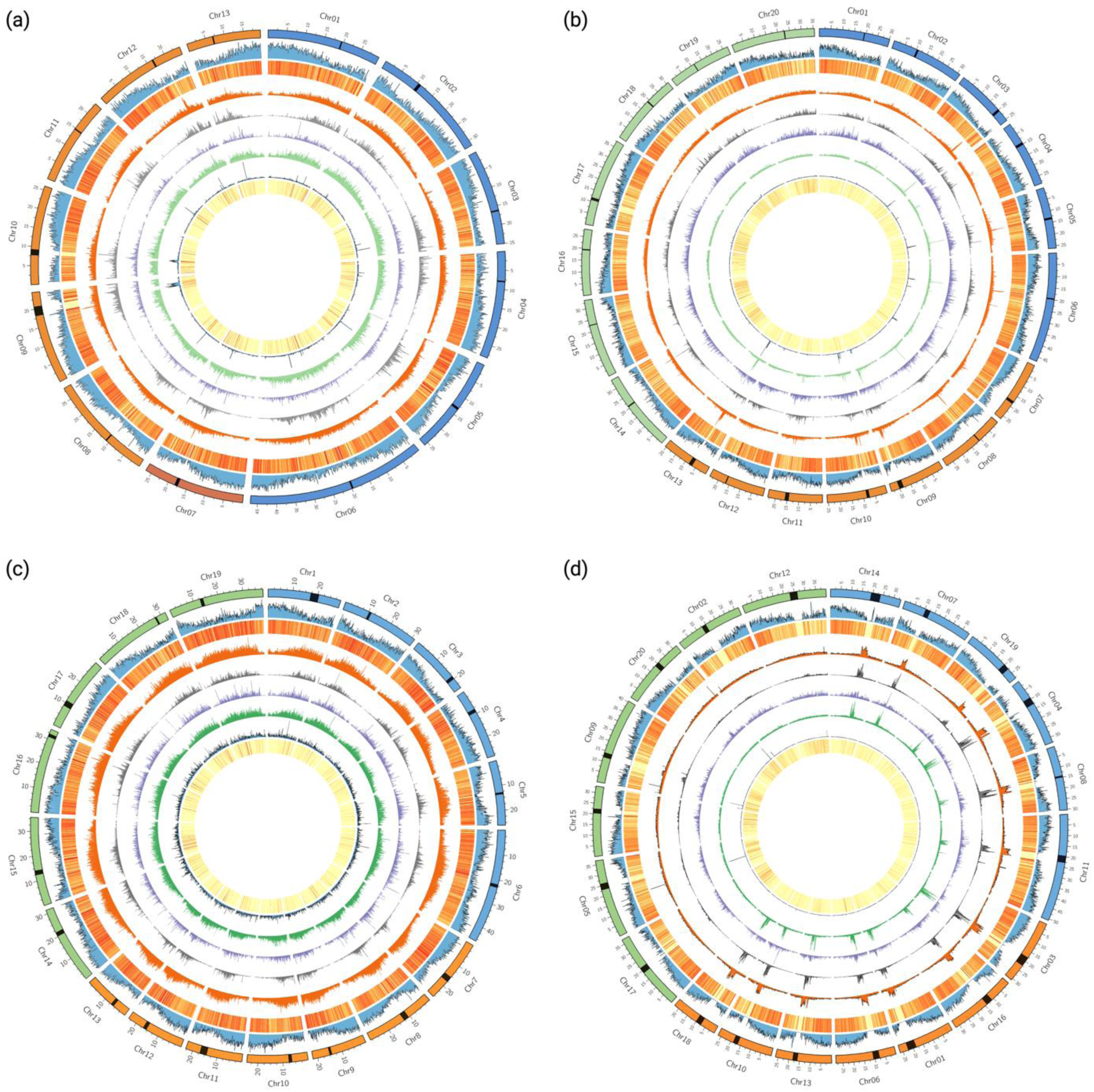
Genome of Camelina species. (a) *Camelina microcarpa* tetraploid Cmi4X, (b) *C. microcarpa* CmiT1, (c) *C. microcarpa* CmiT2, and (d) *C. sativa* CsaDH55. The outer track represents chromosomes and subgenomes by blue, orange, and green colour and centromeres with a black band. The tracks from outer to inner circle represents gene density, expression of genes, repeat distribution, gypsy distribution, copia distribution, Helitron distribution, mutator distribution, and full-length LTR distribution.

**Table 1.** Features of *Camelina microcarpa* and *C. sativa* genomes.

| **Genomes** | **CN119243** | **CN120025** | **CN119205** | **DH55** |
| --- | --- | --- | --- | --- |
| Estimated genome size (Mb) | 419.6 | 577.3 | 649.2 | 711.9 |
| Assembled genome size (Mb) | 384.06 | 555.63 | 607.97 | 678.31 |
| No. of chromosomes (n) | 13 | 19 | 20 | 20 |
| Scaffold N50 (Mb) | 26.09 (7) | 29.88 (8) | 28.42 (9) | 32.12 (10) |
| Scaffold N90 (Mb) | 19.45 (13) | 22.87 (17) | 23.24(19) | 26.37 (19) |
| N counts | 3,500 | 258,100 | 11,200 | 3,200 |
| Number of repeat elements | 230,176 | 322,719 | 403,779 | 616,027 |
| Percentage of repeat elements | 41.36 | 42.40 | 44.87 | 50.25 |
| Total number of full-length LTR | 1,905 | 3,194 | 4,057 | 4,576 |
| Copia (%) | 4.97 | 5.09 | 5.15 | 7.08 |
| Gypsy (%) | 11.80 | 12.93 | 13.03 | 12.39 |
| Helitron (%) | 15.60 | 15.90 | 17.28 | 19.95 |
| other repeats (%) | 8.99 | 8.48 | 9.42 | 10.83 |
| LAI | 22.2 | 22.23 | 22.49 | 18.27 |
| Total number of genes | 65,354 | 95,759 | 101,172 | 102,393 |
| BUSCO (%) | 99.2 | 99.4 | 99.4 | 99.5 |

Repeat elements comprised 41.36%, 42.40%, 44.87%, and 50.25% of the assembled genomes of Cmi4X, CmiT2, CmiT1, and CsaDH55, respectively where Gypsy (11.80-13.03%), Copia (4.97-7.08%) and Helitrons (15.6-19.95%) are the dominant classes of repeat elements (**Supplementary Table 3**). The number of full-length LTR elements as might be expected correlated with the genome size; utilizing these sequences to identify the superfamilies among the Copia and Gypsy elements, a similar pattern of prevalence was found in each of the *Camelina* genomes and sub-genomes, with the notable exception being the levels of Ale (Copia), Athila (Gypsy) and Tekay (Gypsy) elements in SG3 of the hexaploids (**Supplementary Table 4a**). As a proportion of the total elements, the Ale elements decreased somewhat in the third sub-genome while the Tekay elements significantly proliferated (**Supplementary Table 4b, Supplementary Fig. 3**). The increase in Tekay elements in the third subgenome is a recent expansion event (<2mya) and would presumably have contributed to the evolution of the SG3 in each hexaploid. The pattern of repeat expansion is mirrored in the third subgenomes of *C. microcarpa* T1 and *C. sativa*, while the *C. microcarpa* T2 showed a lower percentage of Ale elements and an increase in Tekay elements, it also showed a unique small increase in Athila elements (**Supplementary Fig. 3**). The most obvious feature was the expansion of the total number of full-length LTR elements in the third sub-genome for all three hexploid species, although to a lesser extent in *C. microcarpa* T2, with the average prevalence increasing from one every ∼200 Kb in any of SG1 or SG2, to one every ∼100 Kb in SG3 (**Supplementary Fig. 4**). Based on the centromeric retrotransposon repeats (CRB) reported in *Brassica rapa* (Lim et al., 2007), tentative centromeric positions were assigned with possible pseudo-centromeres identified in chromosomes of the third subgenome (**Supplementary Table 5**). It was noted that there was a proliferation of repeat elements around the centromeres of SG1 and SG2 for CsaDH55, particularly Gypsy elements and Helitrons, this was also observed but to a lesser extent for CmiT1, which share the n=20 karyotype, the same subgenome bias was not found for CmiT2 (**Fig. 1**).

Helixer (Holst et al., 2023), a machine learning based *de novo* prediction tool for gene annotation, was used to predict gene and protein models in the respective genomes (**Supplementary Table 6**). We predicted 65,354 protein coding genes in Cmi4X, 95,759 genes in CmiT2, 101,172 in CmiT1 and 102,393 predicted genes in CsaDH55, with associated high confidence BUSCO scores (**Supplementary Fig. 5**). In the tetraploid and all three hexaploids, SG2 had the lowest gene density while SG3 had the highest. The tetraploid (Cmi4X) genome contained relatively higher numbers of tandemly and proximally duplicated genes compared to the first two subgenomes in all hexaploids, which suggests either loss of such duplicates after the whole genome duplication process or the more recent proliferation of these genes in the tetraploid. There was generally a higher number of tandem and proximal duplicate copies in the third subgenome of the hexaploid *Camelina* species (**Supplementary Table 7**). Gene fractionation was estimated using syntenic gene analysis; across all genomes 22,843 *A. thaliana* syntenic orthologs were present, of these 93.40% were found in diploid *C. neglecta* (Chaudhary *et al*., 2023b), while a reduction was seen to ∼90% in all *C. microcarpa* subgenomes, the third subgenome of *C. sativa* (CsaDH55) appeared to show some level of fractionation with only 86.75% syntenic genes being present, although this was confounded by the increased presence of tandemly repeated genes, which were not necessarily identified by the synteny analyses (**Supplementary Table 8**).

### Evolutionary relationship among *Camelina* species

The evolutionary relationships among the *Camelina* species were studied with different approaches, where the four genomes from this study along with *C. neglecta* (Chaudhary *et al*., 2023b), *C. hispida* (Martin *et al*., 2022), and *C. laxa* (Martin *et al*., 2022) were utilized. First, assembled genomes were compared to observe the collinearity among different *Camelina* species using whole genome alignment (**Supplementary Fig. 6**). The relationship between the lower chromosome number species and the hexaploids was as expected based on previous genotypic and cytogenetic work (Chaudhary *et al*., 2020), yet notably the chromosome structure was incredibly conserved with very limited large structural variation differentiating the genomes. Cluster analysis using Orthofinder v.2.5.2 (Emms and Kelly, 2019) further confirmed the relationships among the species (**Fig. 2a**), with SG1 of *C. sativa* (CsaDH55), *C. microcarpa* (Cmi4X, CmiT1, CmiT2) and *C. neglecta* clustering together; similarly, SG2 of the tetraploid and the three hexaploids were clustered. Interestingly, while SG3 of *C. sativa* (DH55) and *C. microcarpa* T1 (CN119205) showed a close relationship with *C. hispida*, the third subgenome of *C. microcarpa* T2 formed a separate clade relatively close to the *C. neglecta* genome. While the outgroups of *A. thaliana*, *C. laxa*, and *C. hispida* possessed higher numbers of unique orthogroups, SG3 from the *C. microcarpa* T2 hexaploid ranked just below them, and *C. sativa* DH55 possessed the lowest number of species specific orthogroups for all the subgenomes (**Supplementary Table 9**).

**Figure 2.**
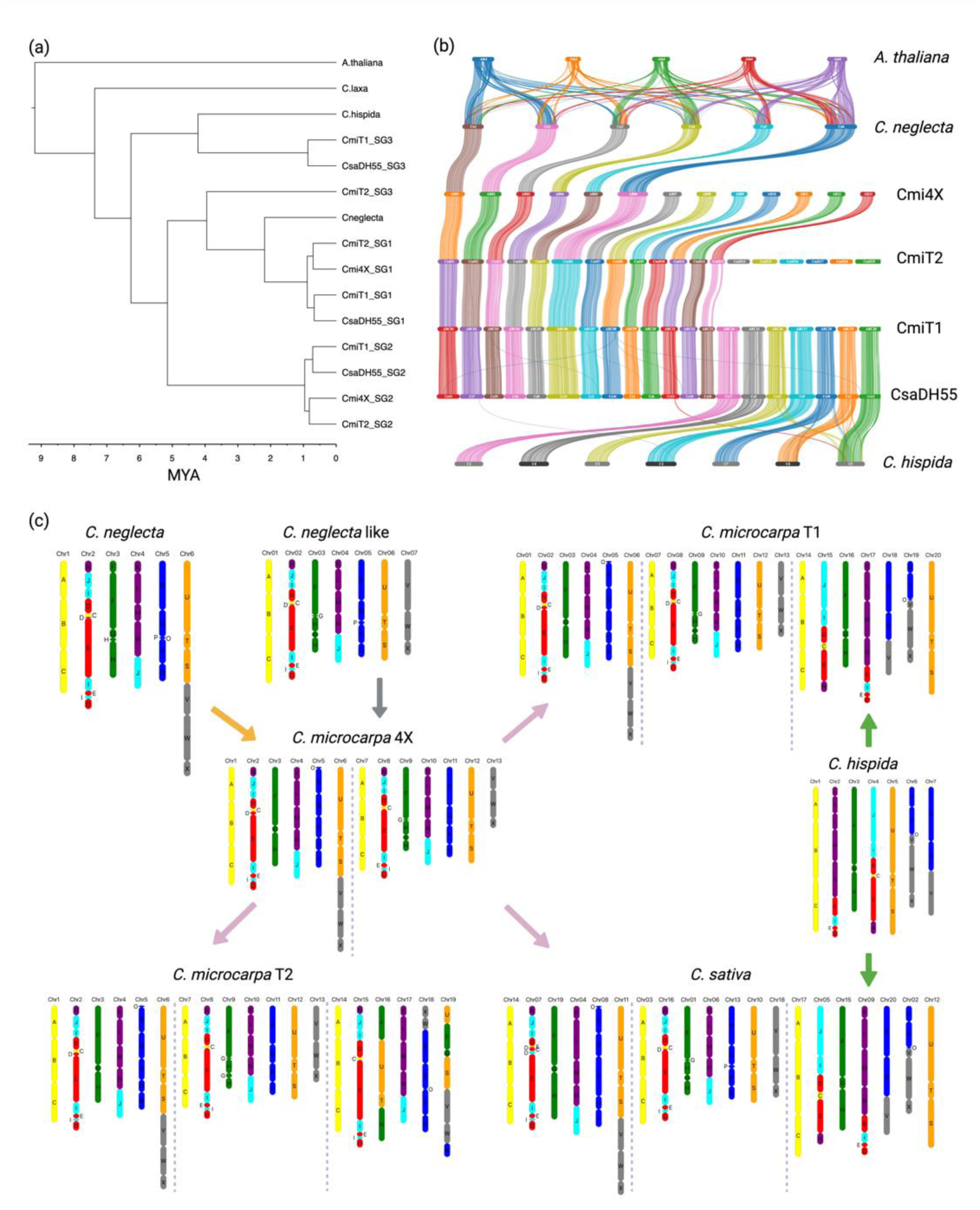
The evolution of *Camelina* species. (a) The species tree represents the relationship among different subgenomes and species of *Camelina* with approximate age of divergence in million years ago (MYA); (b) A syntenic relationship among different *Camelina* species and *Arabidopsis thaliana* showing relationship between subgenomes across Camelina species; and (c) Genomic block arrangements in *Camelina* species and probable evolutionary trajectory across various ploidy levels of *Camelina* species. CmiT1 is *C. microcarpa* Type 1, CmiT2 is *C. microcarpa* T2, Cmi4X is tetraploid *C. microcarpa*, and CsaDH55 is *C. sativa*.

Syntenic analysis was performed using MCScanX (Wang et al., 2012) based on ancestral genomic blocks reported in *A. thaliana* (Schranz et al., 2006); all 24 genomic block (GBs) were identified in all genomes (**Supplementary Table 10**). The visualized syntenic analysis (Bandi and Gutwin, 2020) corroborated the conserved nature of the genome structure among the species, including all rearrangements in SG1 and SG2 relative to the ancestral genome (**Fig. 2b**; **Supplementary Table 10**). The GB arrangement for SG1 of all the assembled higher ploidy genomes is similar to *C. neglecta*, except for chromosome 5 which shows evidence of rearrangement (**Fig. 2c**). *Camelina hispida* showed strong synteny with SG3 of *C. microcarpa* T1 (CmiT1) and *C. sativa* (CsaDH55), which would align with the suggestion that *C. hispida* could be an extant progenitor for SG3 of n=20 *Camelina* species; however, there has been a conserved rearrangement of GBs of chromosome 2 of *C. hispida* relative to the hexaploids (**Fig. 2c**). The origin of the lower chromosome number SG3 of *C. microcarpa* T2 has yet to be found based on the now characterized genomes of the Camelina genus; although this subgenome shows relatively strong similarity with *C. neglecta* and hence SG1 of the other *Camelina* species, significant reshuffling of ancestral GBs has occurred for chromosomes 16, 18 and 19 (**Fig. 2c**). Multiple recombination events would need to be inferred between chromosome 3, 5 and 6 of a *C. neglecta* like genome to result in the final chromosomes 16, 18 and 19 of *C. microcarpa* T2 (**Supplementary Fig. 7**).

The syntenic gene pairs identified among the subgenomes of the different *Camelina* species were utilized to analyze the synonymous substitution rate (Ks) using the GenoDup pipeline (Mao, 2019). As previously reported in Chaudhary *et al*. (2023b), SG1 of all higher ploidy *Camelina* species diverged from *C. neglecta* at approximately the same time, similarly SG2 of all higher ploidy *Camelina* species diverged from *C. neglecta* around the same time (**Fig. 3a**). In the same comparisons with *C. neglecta*, SG3 of CmiT1 and CsaDH55 showed a common slightly older peak compared to that of SG2, which was in contrast to SG3 of CmiT2 where the peak suggested a more recent divergence from *C. neglecta* (**Fig. 3a)**. On comparison with *C. hispida,* the other potential progenitor diploid genome, there was no clear differentiation between the subgenomes of Cmi4X, as well as the first two subgenomes of each of the hexaploids. However, for CmiT2 all three subgenomes shared a peak of divergence, which again differentiated this species from both CmiT1 and CsaDH55, where SG3 showed a more recent divergence event from the progenitor (**Fig. 3b**).

**Figure 3.**
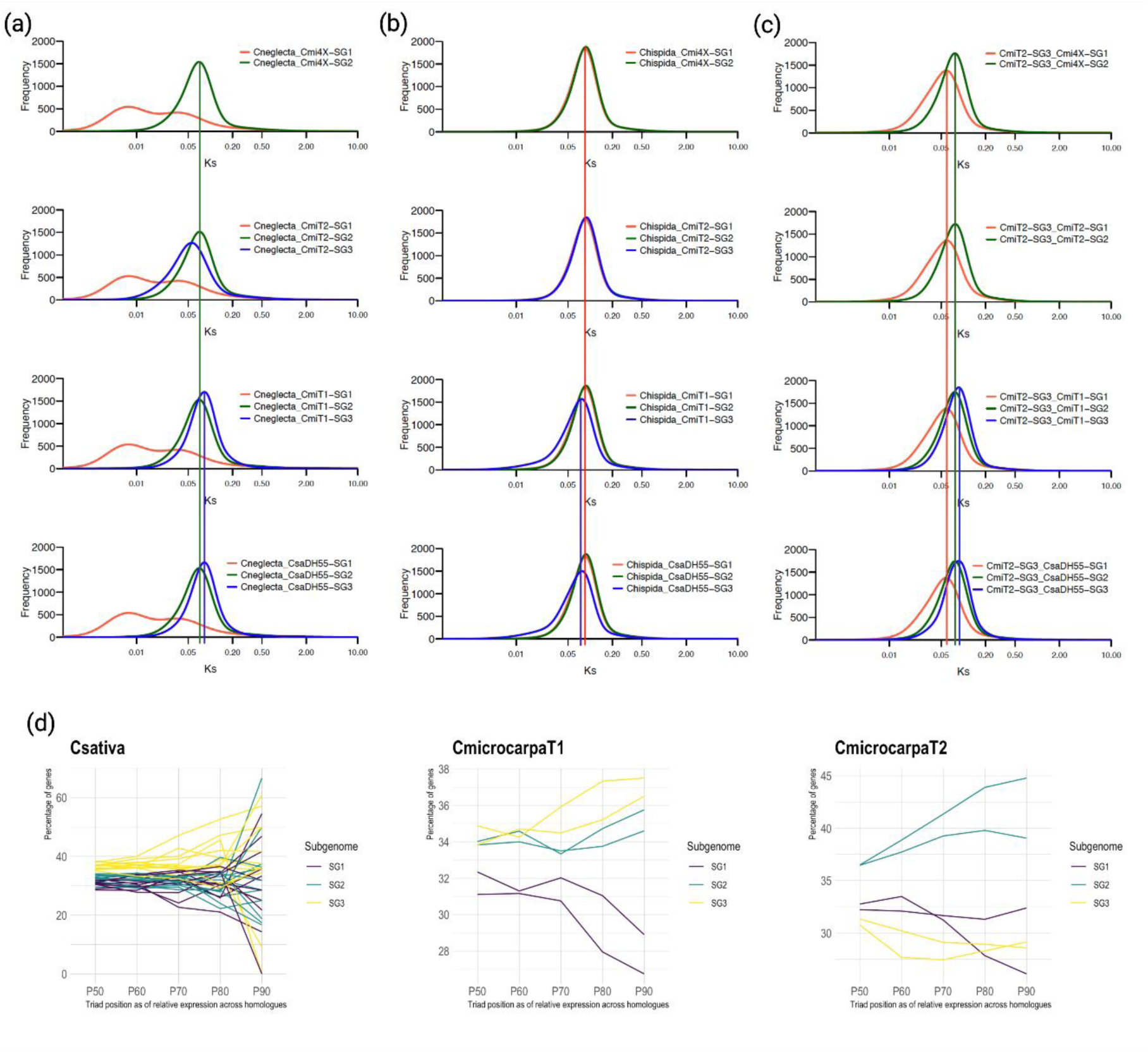
Subgenome evolution in *Camelina* species. Ks distribution of *Camelina neglecta* (a), *C. hispida* (b), and *C. microcarpa* T2 (c) relative to subgenomes of other *Camelina* species, the legend shows the genepairs of the subgenomes from which Ks analysis were performed; and (d) Subgenome dominance analysis in *Camelina* species, the plots shows percentage of genes for particular subgenome (SG1: Purple, SG2: Blue, and SG3: Yellow) with a biased gene expression of at least 50-90% (P60: 60%; P70:70%; P80:80% and P90:90%) in a triad comparison among the subgenome orthologues. *Camelina sativa* shows expression for 12 tissues from *C. sativa* var. DH55 (adopted from Kagale et. al., 2016), *C. microcarpa* T1 shows leaf tissue data for two genotypes (TMP26168 and CN 119205) and *C. microcarpa* T2 show leaf tissue data for two genotypes (CN12025 and CN115248).

Comparing Ks rates among higher ploidy *Camelina* species, identified a barely indistinguishable peak for the first and second subgenomes of all three hexaploids, either when compared with the tetraploid or when compared among the hexaploids (**Supplementary Fig. 8**). However, SG3 of CsaDH55 notably showed two equal peaks when compared with SG3 of CmiT1, but not CmiT2. Although *C. sativa* and CmiT1 appear to share a common progenitor for SG3, the additional peak could suggest that the *C. hispida* like species was added to a tetraploid genome in two different events, leading to separate lineages for *C. sativa* and CmiT1 (**Fig. 3c**).

Subgenome dominance analysis or variance of gene expression (FPKM) was performed with the sets of triplicated orthologous genes (17,378 - 19,278 triplicates) identified among hexaploid *Camelina,* with two genotypes of each species used. The data from *C. sativa* (DH55 and CN120013) were mapped to the CsaDH55 reference genome, while that of *C. microcarpa* T1 (CN119205 and TMP26168) were mapped to the CmiT1 genome reference, and that of n = 19 chromosome *C. microcarpa* T2 data (CN120025 and CN115248) were mapped to the CmiT2 reference. The third subgenome was found to be dominant for all genotypes with *C. sativa* type genome structure, whereas, for *C. microcarpa* T2 (n=19), the second subgenome was found to be dominant over the other two subgenomes (**Fig. 3d, Supplementary Fig. 9, Supplementary Table 15**). These results might indicate that subgenome dominance in *Camelina* species was determined by the nature of the progenitor genome hybridized with the tetraploid structure, rather than the order of events.

The HiC data generated for the assembled genomes, from which physical chromosomal proximity can be inferred, were utilized to assess potential relationships between the subgenomes of the polyploid species. The Iterative Correction of HiC data (ICE) normalized matrices were used to identify the number of significant interactions within and between the chromosomes using fithic2 (Kaul et al., 2020). A total of 684,083 significant interactions (q-value<0.01) were found in the tetraploid Cmi4X, where intra-chromosomal interactions dominated at 90.1% while inter-chromosomal interactions were 9.9%, of which 42.8% were between sub-genomes, and there were slightly less significant interactions among the chromosomes of the second subgenome compared to the first (**Supplementary Fig. 10a; Supplementary Table S12 and S13**). In CmiT2, 88.93% and 11.06% of the 2,090,111 significant interactions were intra- and inter-chromosomal, respectively (**Supplementary Fig.10b; Supplementary Table S12 and 14**). SG1 and SG3 showed the highest number of inter-chromosomal interactions (67.97%) among the sub-genome pairs. In the case of CmiT1, a total of 998,360 significant interactions were found, among these 89.04% are intra-chromosomal and 10.9% are inter-chromosomal (**Supplementary Fig.10c; Supplementary Table S12 and S15**). Among the sub-genome pairs SG1 and SG2 showed the highest number of inter-chromosomal interactions, 37.09% compared to 31.53% between SG1 and SG3 and 31.38% between SG2 and SG3. Interestingly after filtering of a similar amount of HiC data, a smaller number of significant interactions could be detected for CsaDH55; a total of 147,541 interactions were found and among these 90.86% were intra-chromosomal while 9.14% were inter-chromosomal (**Supplementary Fig.10d; Supplementary Table S12 and S16**). Almost all the inter-chromosomal interactions were between homoeologous chromosomes and similar to CmiT1, SG1 and SG2 showed the highest number of significant interactions (36.3%), while SG3 interacted less with both SG1 (33.5%) and SG2 (30.2%). It was evident that homoeologous chromosomes have some level of association and presumably proximity in the nucleus of the polyploids. Three dimensional images which visualize the chromosome positions within the nucleus, effectively simulating their proximity and the interactions among the homoeologous chromosomes in *Camelina* species were generated from chromatic capture information using NucProcess (Stevens et al., 2017) (**Fig. 4**). As might be expected the chromosomes each formed clear domains and the differential interactions between the sub-genomes could be visualized. In order to confirm the observed differences, further filtering of the significant interactions derived from fitHiC was carried out, removing those bins with interactions >1000, which could represent erroneous read mapping to repetitive regions. In addition, additional HiC data generated using an alternative method (Proximo compared to Omni-C) was analysed using the NucProcess method (**Supplementary Table 12**). In all instances CmiT1 and CsaDH55 showed higher levels of interactions between SG1 and SG2 while CmiT2 showed higher levels of interactions between SG1 and SG3.

**Figure 4.**
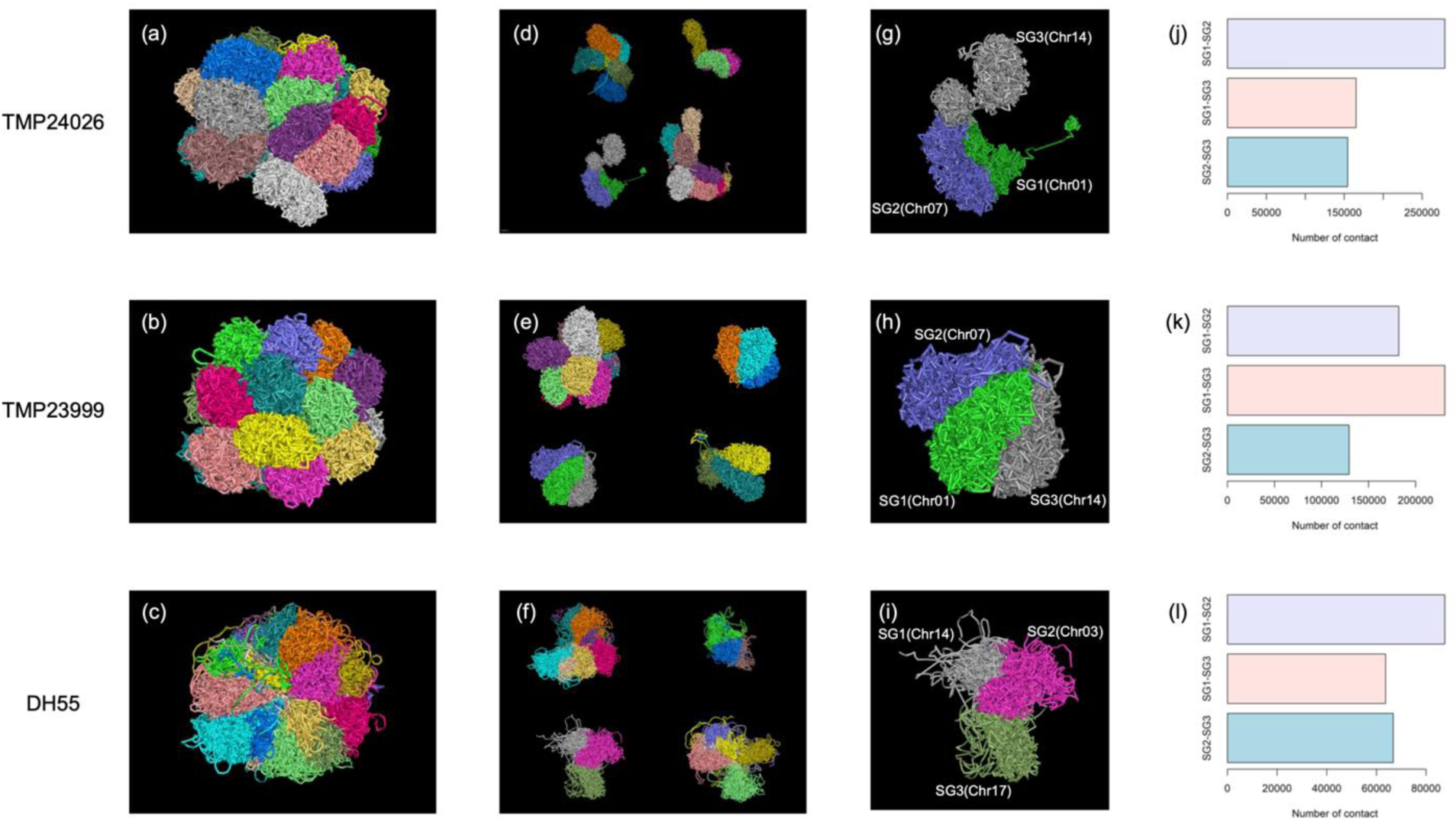
3D structure of *Camelina* species showing interaction among chromatids as well as homoeologous chromosomes. **a-c**. 3D structure of the whole genome; **d-f**. Drag-out of homologous chromosomes; **g-i**. Magnification of three homologous chromosomes; **j-l**. Comparison of number of contacts between different homologous chromosomes. TMP24026 is CmiT1, TMP23999 is CmiT2 and DH55 is CsaDH55 genome.

### Evolution at the gene family level: Resistance Gene Analogs and *Flowering Locus C* in Camelina species

The Resistance Gene Analogs (RGA) were analyzed for all the assembled genomes using RGAugury (Li et al., 2016) (**Supplementary Table 17**). The *C. microcarpa* T1 genome had a slightly higher number of RGA compared to other genomes; however, both hexaploid *C. microcarpa* species possessed higher numbers of RGA clusters, containing at least 10 RGA (**Supplementary Table 18**). Receptor-like kinases (RLK) were the dominant class for all the species. Although there was some loss and gain of RGA in clusters across species, in general the clusters were found in conserved positions across the *Camelina* species. Closer inspection of the largest cluster, observed on chromosome 6 of *C. neglecta,* showed RGA expansion in the *C. microcarpa* species, and presumably a subsequent loss of genes in the derived *C. sativa* genome (**Fig. 5a**). The variable pattern of gene retention was invariably due to loss and gain of RLK loci.

**Figure 5.**
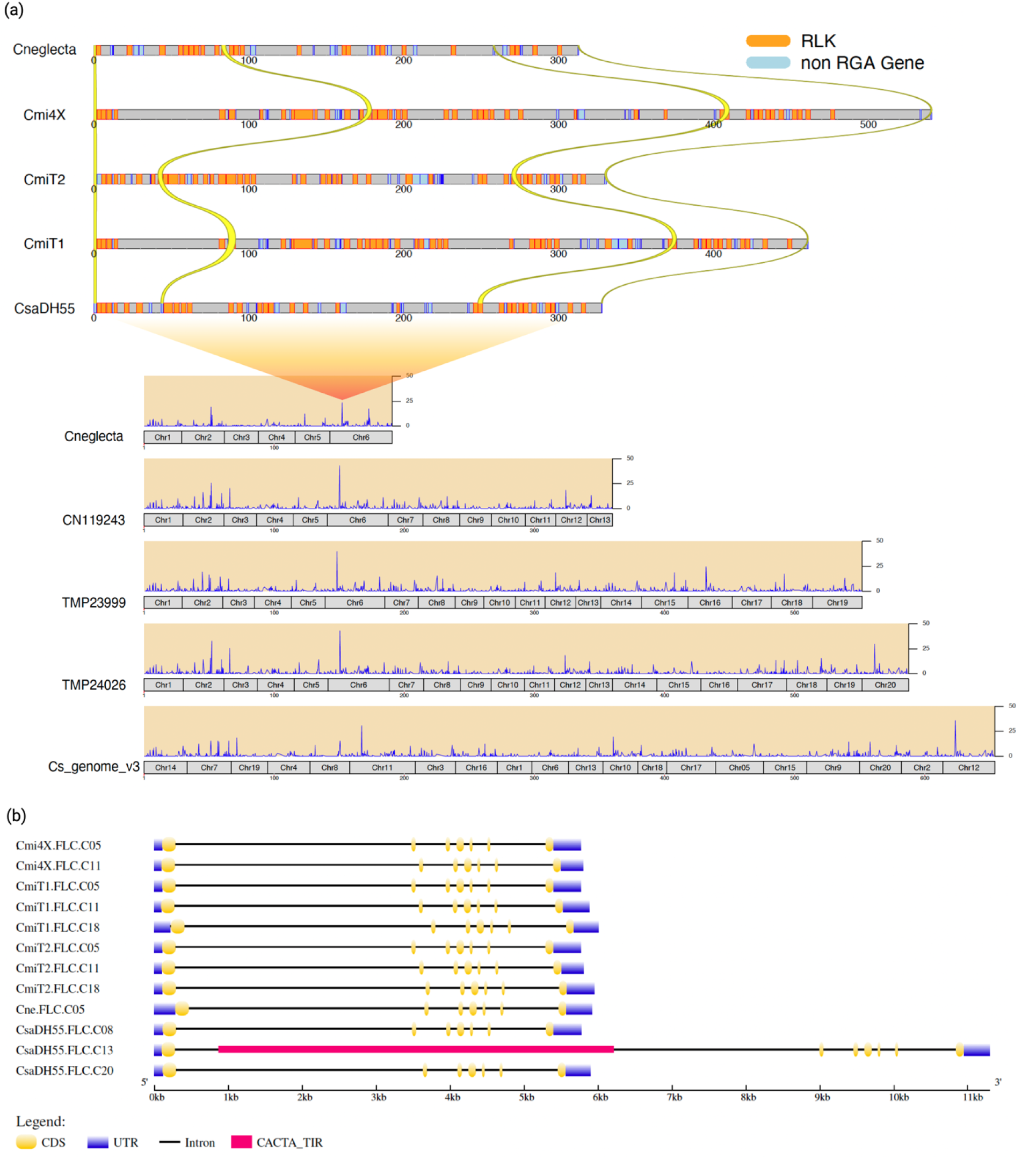
Evolution of resistance genes and flowering gene in *Camelina* species. (a) RGA clustering in *Camelina* species showing conservation of RGA; and (b) Conservation of *FLC* structure in *Camelina* species with a divergent in *Csa.FLC.C13* structure by an insertion of a transposon (red). CmiT1 is a *C. microcarpa* T1; CmiT2 is a *C. microcarpa* T2, Cmi4X is a tetraploid *C. microcarpa*, and CsaDH55 is *C. sativa*.

Due to the importance of *Flowering Locus C* (*FLC*) in determining the growth habit of *C. sativa* (Anderson et al., 2018; Chaudhary et al., 2023a), the gene structure was studied in the assembled Camelina genomes, with *C. sativa* representing the only species with a spring growth habit. As expected, a single copy of *FLC* was found in each subgenome. A number of variants were conserved across species suggesting they did not contribute to phenotypic variance, the exceptions being a deletion of three bases from *Csa.FLC.C8* on chromosome 8 and an insertion of three bases on *Csa.FLC.C20* on chromosome 20, both unique to *C. sativa*. A large structural variation was observed for *Csa.FLC.C13* on chromosome 13 in CsaDH55, where there was an insertion of a 5.3 Kb transposon (CACTA_TIR) in the first intron (**Fig. 5b**). The exact effect of such insertions has not been identified; however, similar events have been reported in *Brassica* species to be associated with the regulation of *FLC* genes (Kitamoto et al., 2014; Yin et al., 2020).

### Potential to exploit the genetic diversity of *Camelina microcarpa* “Type 2”

Genetic diversity among 18 *C. microcarpa* T2 genotypes was analyzed against the CmiT2 reference using genotype by sequencing (GBS) data from Chaudhary *et al*. (2020). The species previously had been noted to have higher genetic diversity compared to *C. sativa* and the principal coordinate analysis showed a higher dissimilarity among these genotypes, with 30.3% of variation on PC1 and 6.96% of variation on PC2, supporting two subpopulations (**Fig. 6a**) (**Supplementary Fig. 11**). The differentiation was also high (F_ST_=0.221) between these two subpopulations (**Supplementary Table 19**).

**Figure 6.**
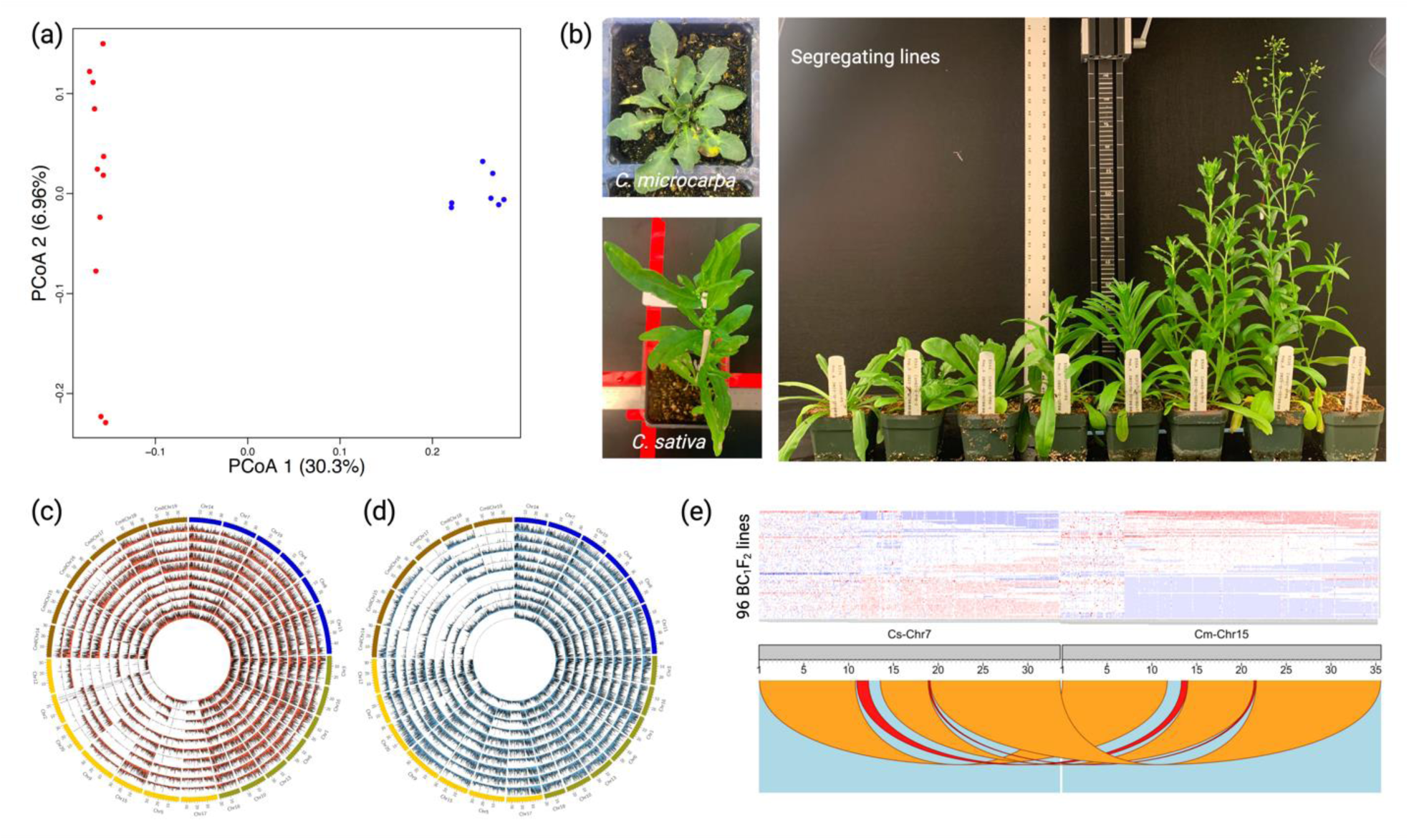
Utilization of *Camelina microcarpa* T2 to increase genetic diversity. (a) Principal Coordinate Analysis among the CmiT2 lines to represent genetic diversity; (b) Leaf morphology of segregating plants developed from interspecific hybridization between *Camelina microcarpa* × *Camelina sativa* ; Distribution of reads mapped to the pseudo-genome coming from segregating lines F_2_-83-8 (c), F_2_-82-6 (d), the blue, green and yellow bands in the outer track represent SG1, SG2 and SG3 from CsaDH55, and the brown bands represent SG3 of CmiT2, inner individual tracks represents reads from different F_2_ lines mapped to the pseudo-genome; and (e) homoeologous recombination in 96 segregating lines as represented by loss/gain of reads represented by blue/red colour on top, individual rows represent different segregating lines, the syntenic relationship between homoeologous chromosomes are represented by links at the bottom of the image.

In order to assess the feasibility of capturing some of the novel diversity from the wild relative, hybridization of CmiT2 with *C. sativa* was performed. A total of 49 F_2_ plants were obtained from four F_1_ progeny from a cross between *C. microcarpa* T2 (CN119102) and *C. sativa* (CN120019), of which four plants showed spring-type early seedling growth, while the remainder of the plants were winter-type. There was a marked difference in leaf shape, leaf waxiness and number of flowers in the segregating plants in comparison to the parents (**Fig. 6b**). Among the 49 plants, only 7 were completely fertile, 3 were partially sterile and the remainder were sterile (**Supplementary Table 20**). It was anticipated that homologous chromosome pairing, and recombination would occur between the common subgenomes 1 and 2, but the disparity between the third subgenomes, which differentiates the parental lines, would result in some abnormal chromosome pairing. Genetic analysis of the F_2_ lines using GBS data, where sequence reads were mapped to a pseudo-genome containing all four sub-genomes (see methods), indicated that SG1 and SG2 did pair relatively normally, but a non-random mapping of sequence reads was observed for SG3. The segregating plants showed a bias towards retention of either SG3 from *C. sativa* or CmiT2, according to the parental hybrid from which the progeny were derived (**Fig. 6c & 6d**), this bias was also reflected in the leaf morphology of the progeny (**Supplementary Fig. 12**). A number of the plants showed missing chromosomal segments either due to deletion or translocation, probably resulting from homoeologous recombination.

In order to further assess recombination events between *C. sativa* (CN120019) and *C. microcarpa* (CN119102), one of the hybrids was backcrossed with PI650132 (a spring-type *C. alyssum*, considered a cryptic species of *C. sativa*), the resulting backcross population was self-pollinated to generate BC_1_F_2_ seeds, of these 96 lines were genotyped using GBS (**Fig. 6e**). Mapping of sequence reads from the 96 individuals to the pseudo-genome identified a similar overall pattern with biased retention of the third subgenome of the *C. sativa* type. Studying some of the homoeologous regions, there was evidence of exchange; a uniform distribution of mapped reads was observed across chromosome 15 of CmiT2, yet based on read depth, the reads segregated for the absence (AA), single copy presence (AB) and presence (BB), of homoeologous regions of chromosome 15 and chromosome 7 of CsaDH55, respectively, suggesting a homoeologous recombination event between these chromosomes (**Fig. 6f**). Beside this event, some of the lines were found to be missing whole chromosomes such as chromosome 17 and chromosome 2 of *C. sativa*.

## Discussion

Genome assemblies and associated resources are important tools in the study of polyploids and identifying relationships among closely related species can reveal distinguishing genomic features, which shape their genome evolution. Previously, genome assemblies of two diploid *Camelina* species had shed light on the evolutionary history of *C. sativa; C. neglecta* was identified as a progenitor of the first subgenome of hexaploid *C. sativa*, and it had been suggested that *C. hispida* was an extant progenitor of the third subgenome (Chaudhary *et al*., 2020; Mandáková et al., 2019). In the current study three new high quality genome assemblies of *C. microcarpa*, of which one is a tetraploid and the other two are hexaploids differentiated by their chromosome number, has provided further insights into the evolution of the *Camelina* genus. The tetraploid *C. microcarpa* genome, which coincides with the first and second subgenome of *C. sativa* and both *C. microcarpa* hexaploids, suggested a step-wise merger of species from diploid (n = 6) to tetraploid (n = 13) and then hexaploid (n=19 and n = 20) (Brock et al., 2022; Chaudhary *et al*., 2023b). Yet the third subgenome of *C. microcarpa* T2, not withstanding the reduction in chromsome number from the common n=7 to n=6, is quite distinct in structure. Although the third subgenome of *C. microcarpa* T2 shared the same number of chromosomes with *C. neglecta,* a number of significant chromosomal rearrangements distinguish the genomes (**Fig. 2**). The Ks analyses suggests a more recent divergence from a *C. neglecta-*like genome compared to SG3 of the other *Camelina* hexaploids, and as yet no extant relative has been collected that resembles this subgenome structure.

The third subgenome of all hexaploids displayed a number of differences, which might relate to their later addition in the evolutionary trajectory of *Camelina* species. There was a higher gene density and the notable proliferation of full-length LTR in the third subgenome; the age of divergence of tandem duplicate genes indicated the third subgenome was added last to the tetraploid like species on the path to the hexaploids (**Supplementary Fig. 13**). Although the rate of synonymous substitutions indicated that the third subgenome from *C. microcarpa* T2 was more recently diverged from *C. neglecta* compared to the second subgenome, the pattern of increased tandem duplicates in the third subgenome also reflected that seen in *C. microcarpa* T1 and CsaDH55. It was proposed that *C. sativa* evolved from a *C. microcarpa* T1 like genome (Brock *et al*., 2018; Mandáková *et al*., 2019), this could not be reconciled with the observation that the third subgenome from *C. sativa* and the third subgenome of *C. microcarpa* T1 showed multiple peaks of divergence. These data suggest two separate lineages, where a *C. hispida* like genome hybridised at different time intervals with the tetraploid structure to form the n=20 chromosome *Camelina* species. A chloroplast based phylogenetic tree derived from the genome data also suggested different potential progenitors for the formation of *C. sativa* and *C. microcarpa* T1 (**Supplementary Fig. 14**). Thus, we speculate although *C. sativa* and *C. microcarpa* T1 share subgenome structures they represent independent evolutionary events.

It is interesting to note that to be in agreement with the “two-step-hypothesis” for the formation of a hexaploid, similar to that inferred in the evolution of related Brassica paleohexaploids (Lyons *et al*., 2008), the last genome added, SG3, should show dominant gene expression. In the case of *C. sativa* and *C. microcarpa* T1, which shared the same subgenome structure, this was true with the third subgenome showing the expected expression dominance. However, in the case of *C. microcarpa* T2, the second subgenome was found to be dominant. The mechanism of dominance has been studied in a number of species and suggested to be associated with a number of genome features within the diploid progenitors, including genome compaction, distribution of repetitive elements and chromosome architecture (Hu et al., 2024). In *Camelina,* there was no such obvious correlation, since the third subgenome for all hexaploids was similar in basic organisation with a higher gene density and a higher number of recently-proliferated full-length LTR elements. The most notable association with the observed genome dominance was that of biased inter-chromosomal interactions for the subgenomes in the hexaploids. Chromosomal 3D architecture has been consistently shown to impact gene expression (He et al., 2024) and the dominant genome, either SG2 in *C. microcarpa* T2 and SG3 for *C. microcarpa* T1 and *C. sativa*, had the least number of significant interactions with their fellow subgenomes, suggesting proximity in the nucleus could be dictating gene expression. It has been shown in a number of species that proximity, as determined through chromosome capture, suggests co-localised homoeologous genes share common expression patterns, which is also reflected in other measures such as methylation status and histone modification patterns (Wang et al., 2021; Zhang et al., 2024). This would be in concurrence with the two non-dominant sub-genomes showing limited evidence of expression bias. Interestingly the alignment of chromosomal proximity with genome dominance has been suggested from work in other allopolyploid species. In cotton it was noted that homoeologous gene pairs with extreme expression bias were found to show fewer chromatin interactions (Wang et al., 2018). In wheat, when comparing gene expression of syntenic triads from the A, B and D genome across multiple tissues, there was shown to be a subtle but consistent bias to the D genome (Ramírez-González et al., 2018) and in Concia et al. (2020) it was found that the A and B genomes interact more frequently than any of the other possibile pairings. Thus it appears that genome-wide expression dominance could result from the higher-order organisation of the chromatin in the nucleus.

The identification of the unique species of hexaploid *C. microcarpa,* Type 2, further the prospects of increasing genetic diversity in the *C. sativa* which has a very low genetic base (Chaudhary *et al*., 2020; Luo *et al*., 2019; Singh *et al*., 2015). Although the clear differentiation of genome structure does raise the issue of likely chromosomal imbalance and thus reduced fertility when trying to cross the two species, which was almost certainly the reason for the anomalies observed in previous studies, which attempted interspecific hybridization prior to the identification of two *C. microcarpa* species (Martin et al., 2019; Tepfer et al., 2020). Yet despite this, as shown here homoeologous recombination with *C. sativa* can result in the capture of genetic diversity from *C. microcarpa* T2. The much higher levels of genetic variation and potential phenotypic variation for traits of interest in *C. microcarpa* T2 suggest further efforts to encourage homoeologous recombination and stabilise the progeny are worth investigation. As knowledge of factors limiting homoeologous recombination in other related species is uncovered (Higgins et al., 2020; Wang et al., 2024), new strategies to affect this work may become feasible.

## Material and methods

### Plant material for DNA extraction and sequencing and assembly

For *C. microcarpa* (accession: CN120025), fresh leaf tissue was flash frozen in liquid nitrogen and stored at -80 °C before DNA extraction. Leaf materials were sent to Dovetail Genomics (Scotts Valley, CA, USA); high molecular weight DNA extraction and long read sequencing using PacBio followed by HiC scaffolding were performed according to the methodology described in Chaudhary et al. (2023b). For the accessions CN119243, CN119205, and DH55 nuclei extraction was done as described by Zhang et al. (2012) followed by DNA extraction using the Nanobind Plant Nuclei Big DNA kit according to the manufacturer’s instructions (Circulomics, Menlo Park, CA, USA). DNA was size selected for >5kb on a BluePippinTM (Sage Science Inc., Beverly, MA, USA) and Oxford Nanopore Technology (ONT) libraries were prepared using Ligation Sequencing Kit V14 (SQK-LSK114) and sequenced with R10.4.1 MinION flow cells on a GridION and MinION platforms from Oxford nanopore Technology. High molecular weight DNA for the same sample were also extracted using NucleoBond® HMW DNA kit (Macherey-Nagel) and sequenced on a PacBio Sequel IIe sequencing platform at the Global Institute for Food Security, Saskatoon, Canada (**Supplementary Table 1**). Illumina short reads (paired end 2 × 125bp) for three accessions (CN119243, CN120025, and CN119205) were generated using the HiSeq2000 Illumina platform (Illumina, San Diego, CA, USA). HiC libraries were prepared using the Dovetail® Omni-C® kit (Cantata Bio, Scotts Valley, California, USA) and the Proximo^TM^ Hi-C kit (Phase Genomics, Seattle, WA, USA) according to the manufacturers protocol; the libraries were sequenced on the Illumina NovaSeq6000 platform (paired end 2 × 150 bp).

Paired Illumina reads were used to estimate the genome size based on k-mer analysis using Jellyfish (Marçais and Kingsford, 2011) followed by visualization using GenomeScope (Vurture et al., 2017). The reads were filtered against the *C. sativa* chloroplast genome (Accession: NC_029337) (Hohmann et al., 2015) using Bowtie2 (Langmead and Salzberg, 2012) before processing for genome size estimation. In the case of DH55, paired raw reads (SRR1171872 and SRR1171873) from Kagale *et al*. (2014) were used. The hexaploid CN120025 was assembled using ∼4 million CCS reads (∼32.9× coverage clean reads) as described in Chaudhary *et al*. (2023b). For the tetraploid (CN119243) and hexaploids (CN119205 and DH55), HiFi data along with ONT data were used to assembled the genome using Hifiasm v0.19.5 with default options (Cheng et al., 2021) (**Table 1**). HiC data generated using omni-C protocol was mapped to the base assembly using HiC-Pro (Servant et al., 2015) and scaffolding was performed using the pipeline EndHiC (Wang et al., 2022), the hic files generated by 3D-DNA (Dudchenko et al., 2017) were visualized using Juicebox (Robinson et al., 2018), manual correction of misassembled regions were performed. Using the fitHiC tool, inter and intra-chromosomal interactions were calculated as described by Zhang *et al*. (2024), except 100 kb bins were used to count the significant interactions.

### RNA extraction, sequencing, and sequence analysis

RNA extraction was performed for two CmiT2 genotypes (CN120025 and CN115248), two *C. sativa* genotypes (DH55, CN120013), two CmiT1 genotypes (CN119205 and TMP26168) and CN119243 (n = 13 chromosomes) with three replications for each. Seeds were grown in CYG seed germination pouches (Mega International, Newport, MN, USA) at room temperature for one week before total RNA extraction from leaf tissue for all samples except CN119243, TMP26168, and CN115248 which were sampled from greenhouse. Total RNA was extracted using a standard RNeasy Plant Mini Kit (Qiagen) as described by the manufacturer with on-column DNA digestion. RNAseq libraries were constructed using the TruSeq RNA preparation kit (Illumina) with 100 ng of RNA for each sample followed by sequencing on an Illumina HiSeq2000 platform (2 × 125 bp). Raw reads were filtered for low quality reads, short reads and adapter contamination using Trimmomatic v.0.33 (Bolger et al., 2014). All trimmed reads were aligned with respective reference genome using STAR v.2.7.6a (Dobin et al., 2013) using default parameters, except for *--alignIntronMax* set at 10000 and *--outFilterMismatchNmax* set at 4. Normalization of read counts was done using the Fragment Per Kilobase of transcripts per Million mapped reads (FPKM) method in RSEM v.1.3.3 (Li and Dewey, 2011).

### Annotation of repetitive elements, gene prediction and centromeric repeat prediction

Helixer (Holst *et al*., 2023) was used for the annotation of all the genomes using model *land_plant*. Repeat elements were annotated using an Extensive de-novo TE annotator pipeline v.2.1.0 (Ou et al., 2019) which uses LTRharvest (Ellinghaus et al., 2008), LTR_finder (Xu and Wang, 2007), LTR_retriever (Ou and Jiang, 2018), TIR_learner (Su et al., 2019), Generic repeat finder (Shi and Liang, 2019) and HeltronScanner (Xiong et al., 2014) to annotate the repeat elements. TEsorter (Zhang et al., 2019) was used to classify the LTR families based on the output from EDTA. Centromeric repeats were predicted as described in Chaudhary *et al*. (2023b).

### Assembly quality assessment

The quality of each genome was assessed using BUSCO v.5.4.3 (Simão *et al*., 2015) analysis utilizing *embryophyta_odb10* (n=1,614) to compare the completeness of single copy orthologs. Also, the quality of the genome assembly was tested by mapping whole genome Illumina short reads to the assemblies using BWA (Li and Durbin, 2009) and the mapping quality was assessed using Qualimap v.2.2.2 (Okonechnikov *et al*., 2016). LTR assembly index (LAI) (Ou *et al*., 2018) was calculated to assess the level of full length LTR repeats as a further measure of genome quality.

### Syntenic analysis

Syntenic analysis was performed to compare gene contiguity among species, using *A. thaliana*, *C. neglecta*, *C. microcarpa* T1, *C. microcarpa* T2, *C. sativa*, and *C. hispida* (Martin *et al*., 2022). Protein models predicted from all three assembled genome were reciprocally blasted against *A. thaliana* to define gene pairs and DAGChainer (Haas et al., 2004) was used to identify syntenic blocks which were assigned to an Cruciferae ancestral genomic block as defined by Schranz *et al*. (2006). Custom scripts were used to draw karyotype blocks for all the assembled genomes and the syntenic genes were visualized using the online platform SynVisio (Bandi and Gutwin, 2020).

### RGA, orthogroups and duplicate gene analysis

The protein models for each genome were used to predict resistance gene analogs (RGAs) using the *RGAugury* pipeline v. 2.1.3 (Li *et al*., 2016). RGA gene clusters were identified as described in Bayer et al. (2019). Protein models were used for orthogroup identification using Orthofinder v.2.5.2 (Emms and Kelly, 2019), orthogroups were compared among *Camelina* species viz. *C. microcarpa* T1, *C. microcarpa* T2, *C. hispida*, *C. laxa*, *C. neglecta* and *C. sativa,* and finally duplicated genes were identified using DupGen_finder pipeline (Qiao et al., 2019), which identified tandem duplicates, proximal duplicates, transposed and whole genome duplicate gene pairs in assembled genomes.

### Estimation of genomic age of divergence

*C. neglecta* (Chaudhary *et al*., 2023b) and *C. hispida* (Martin *et al*., 2022) were utilized to predict the age of divergence among different subgenomes of *Camelina* species. The analysis used syntenic gene-pairs among different species of *Camelina,* rate of synonymous substitutions (Ks value) for each gene-pair was calculated using GenoDup pipeline (Mao, 2019), which uses TranslatorX (Abascal et al., 2010) and MAFFT (Katoh and Standley, 2013) for multiple sequence alignment, and PAML (Yang, 1997) for calculating rate of synonymous substitution between gene-pairs. The Gaussian mixture model from R package mclust (Scrucca et al., 2016) was used to plot distribution of Ks and identify the number of gaussian components. Based on these components, the mean value of Ks distribution was identified and used to calculate the age of divergence as described by Kagale *et al*. (2014). Further, MCMCtree (Yang, 2007) was used to infer the age of divergence from the single copy orthologs from subgenomes of *Camelina* species and *Arabidopsis thaliana;* where the divergence of *A. thaliana* with *Camelina* species was assumed as 9.9 million years ago (Kumar et al., 2022).

### Subgenome dominance analysis

Subgenome dominance was analyzed for two genotypes of each of the hexaploids *C. sativa, C. microcarpa* T1 and *C. microcarpa* T2. Based on syntenic analysis with *A. thaliana,* orthologues in the subgenomes were identified for comparison. Genes expressed at less than 0.01 FPKM in any of the replications or for any of the orthologues were discarded. Using three replications, for each set of orthologues genes in the hexaploid, analysis of variance was performed with a custom script in R statistical software v.4.1.1 (Team, 2021) to observe differences in expression patterns between the subgenomes. More than 11,000 triplicates were studied for all the samples belonging to hexaploid *Camelina* species, and the difference in the number of genes representing each category was determined using an analysis of variance test with a significant threshold of *P-value* < 0.05. Further subgenome dominance analysis were also performed by comparing relative expression of three homoeologues (Ramírez-González *et al*., 2018), where the percentage of genes representing triad position above 50% are counted for all three subgenomes to plot the percentage of gene showing bias in a triads (**Supplementary Fig. 9**).

### 3D model construction for the whole genome structure

Raw Fastq files from HiC experiments were processed using the NucProcess tool (Stevens et al., 2017) to create HiC contact files. The output is generated in Nuc Chromatin Contact (NCC) format. Random 500,000 valid contacts were selected three times for each sample to create 3D coordinates in PDB (ProteinDataBank) format using NucDynamics tool (https://github.com/tjs23/nuc_dynamics). The 3D genome structure was viewed and expanded using PyMOL (Version 3.0 Schrödinger, LLC). The syntenic genomic blocks among three subgenomes were extracted for contact number comparison. Only blocks with similar genomics lengths (no more than 30% length difference among three syntenic blocks) were selected for valid contact extractions to avoid the potential bias caused by the different genome fragment sizes of syntenic blocks. Contact numbers between SG1 and SG2, SG1 and SG3, or SG2 and SG3 were calculated separately for the comparison in different accessions.

### Genetic diversity analysis among *Camelina microcarpa* Type 2 accessions

For genetic diversity analysis, the genotyping by sequencing (GBS) data from Chaudhary *et al*. (2020) was used where the genotypes with 19 chromosomes were selected. Read mapping were performed against the CmiT2 assembly and SNP calling were performed as described in Chaudhary *et al*. (2020). GenAlEx v.6.5 (Peakall and Smouse, 2012) was used to determine the heterozygosity and population differentiation, STRUCTURE (Pritchard et al., 2000) with a burn-in period of 150,000 steps and 150,000 MCMC was used to determine the admixture in the genotypes and the number of subpopulations, AveDissR Package in R (Yang and Fu, 2017) was used for principal component analysis and MEGA X (Kumar et al., 2018) was used to draw the Neighbor-Joining tree.

### Development of interspecific segregating populations

Interspecific hybridization between *C. sativa* (CN120019) × *C. microcarpa* (CN120025) was performed and a limited number of seeds were obtained with *C. sativa* as the maternal parent. In these crosses, only 6 pods (1 seed) formed after manual pollination of 190 flowers. The reciprocal cross, with a different accession of *C. microcarpa* (CN119102), produced 71 seeds from 13 pods. Most of the hybrid seed obtained from these crosses were deformed and only 6 out of 71 seeds germinated. *Camelina microcarpa* was winter-type and required vernalization to reach the reproductive phase whereas CN120019 (*C. sativa*) was a spring-type. The hybrids from these crosses produced a winter-type plant that had intermediate plant and leaf morphology from the parents. The hybrid from the CN119102 × CN120019 cross produced four F_1_ plants, although partially sterile, together they produced 49 F_2_ seeds.

One of the hybrids was backcrossed with accession PI650132, a spring-type *C. alyssum* to generate BC_1_F_1_ plants. A total of 8 seeds were obtained from the (*C. microcarpa* × *C. sativa*) × *C. alyssum* cross; however, only one plant was used to generate the BC_1_F_2_ generations. BC_1_F_2_ plants derived from selfed-seed from the (CN119102 × CN120019) × PI650132 cross were used for genetic analysis.

### Genotyping of segregating lines

One week old leaf tissue was harvested and kept at -80 °C until DNA extraction. DNA extractions were performed using the CTAB method and GBS library preparation was as described by Poland et al. (2012). Paired-end sequencing was done on the Illumina HiSeq platform. Sequences were trimmed for low base quality and adapters using Trimmomatic v.0.33 (Bolger et al., 2014). Reads were mapped to a pseudo-genome generated from combining the CsaDH55 genome with the third subgenome from CmiT2, making a pseudogenome with four subgenome (26 chromosomes), using BWA (Li and Durbin, 2009) with default parameters. SNPs were called using GATK (McKenna et al., 2010) with default parameters. Sequence reads were used to study possible aneuploidy and homoeologous recombination. BEDTools (Quinlan and Hall, 2010) was used to extract mapped reads and to calculate the frequency of mapped reads along 100 Kb bins in the genome. Reads were plotted across the bins to confirm possible genetic events. Circos (Krzywinski et al., 2009) and karyotypeR (Gel and Serra, 2017) were used to plot the distribution of mapped reads along the *Camelina* pseudogenome for visualizing aneuploidy and homoeologous recombination events.

### Chloroplast assembly and analysis

Chloroplast genomes of CsaDH55, CmiT1, Cmi4X, and four *C. sativa* genotypes (MAGIC05, MAGIC08, MAGIC10 and MAGIC15) were assembled from PacBio long read data using Oatk (Zhou et al., 2024), whereas the chloroplast for CmiT2 was assembled using GetOrganelle (Jin et al., 2020) using short Illumina reads. The chloroplast assembly for *Camelina neglecta*, *C. laxa*, *C. hispida* were adopted from (Martin et al., 2022). All the assembled chloroplast were aligned using clustal omega (Sievers et al., 2011) and visualized using MEGA11 (Tamura et al., 2021).

## Data availability

All sequence data is deposited at the EBI-ENA under accession: PRJEB96055. The genome assemblies are available at https://cruciferseq.ca

## References

Abascal, F., Zardoya, R., and Telford, M.J. (2010). TranslatorX: multiple alignment of nucleotide sequences guided by amino acid translations. Nucleic Acids Res 38:W7–13. 10.1093/nar/gkq291.

Al-Shehbaz, I.A., Beilstein, M.A., and Kellogg, E.A. (2006). Systematics and phylogeny of the *Brassicaceae (*Cruciferae): An overview. Plant Systematics and Evolution 259:89–120. 10.1007/s00606-006-0415-z.

Anderson, J.V., Horvath, D.P., Doğramaci, M., Dorn, K.M., Chao, W.S., Watkin, E.E., Hernandez, A.G., Marks, M.D., and Gesch, R. (2018). Expression of *FLOWERING LOCUS C* and a frameshift mutation of this gene on chromosome 20 differentiate a summer and winter annual biotype of Camelina sativa. Plant Direct 2:e00060.

Bandi, V., and Gutwin, C. (2020). Interactive exploration of genomic conservation. Proceedings of the 46th graphics interface conference on proceedings of graphics interface 2020 (GI’20). Canadian Human-Computer Communications Society Waterloo, Canada.

Bayer, P.E., Golicz, A.A., Tirnaz, S., Chan, C.K.K., Edwards, D., and Batley, J. (2019). Variation in abundance of predicted resistance genes in the *Brassica oleracea* pangenome. Plant Biotechnology Journal 17:789–800.

Berti, M., Gesch, R., Eynck, C., Anderson, J., and Cermak, S. (2016). Camelina uses, genetics, genomics, production, and management. Industrial Crops and Products 94:690–710. 10.1016/j.indcrop.2016.09.034.

Bird, K.A., Niederhuth, C., Ou, S., Gehan, M., Pires, J.C., Xiong, Z., VanBuren, R., and Edger, P.P. (2019). Replaying the evolutionary tape to investigate subgenome dominance in allopolyploid *Brassica napus*. BioRxiv:814491.

Bolger, A.M., Lohse, M., and Usadel, B. (2014). Trimmomatic: A flexible trimmer for Illumina sequence data. Bioinformatics 30:2114–2120. 10.1093/bioinformatics/btu170.

Brock, J.R., Donmez, A.A., Beilstein, M.A., and Olsen, K.M. (2018). Phylogenetics of *Camelina* Crantz. (Brassicaceae) and insights on the origin of gold-of-pleasure (*Camelina sativa*). Molecular Phylogenetics and Evolution 127:834–842. 10.1016/j.ympev.2018.06.031.

Brock, J.R., Scott, T., Lee, A.Y., Mosyakin, S.L., and Olsen, K.M. (2020). Interactions between genetics and environment shape *Camelina* seed oil composition. BMC plant biology 20:1–15.

Brock, J.R., Mandáková, T., McKain, M., Lysak, M.A., and Olsen, K.M. (2022). Chloroplast phylogenomics in *Camelina* (Brassicaceae) reveals multiple origins of polyploid species and the maternal lineage of *C. sativa*. Horticulture Research.

Chaudhary, R., Higgins, E.E., Eynck, C., Sharpe, A.G., and Parkin, I.A. (2023a). Mapping QTL for vernalization requirement identified adaptive divergence of the candidate gene *Flowering Locus C* in polyploid *Camelina sativa*. The Plant Genome 16:e20397.

Chaudhary, R., Koh, C.S., Perumal, S., Jin, L., Higgins, E.E., Kagale, S., Smith, M.A., Sharpe, A.G., and Parkin, I.A. (2023b). Sequencing of *Camelina neglecta*, a diploid progenitor of the hexaploid oilseed *Camelina sativa*. Plant Biotechnology Journal 21:521–535.

Chaudhary, R., Koh, C.S., Kagale, S., Tang, L., Wu, S.W., Lv, Z., Mason, A.S., Sharpe, A.G., Diederichsen, A., and Parkin, I.A. (2020). Assessing Diversity in the *Camelina* Genus Provides Insights into the Genome Structure of *Camelina sativa*. G3: Genes, Genomes, Genetics 10:1297–1308.

Cheng, F., Sun, C., Wu, J., Schnable, J., Woodhouse, M.R., Liang, J., Cai, C., Freeling, M., and Wang, X. (2016). Epigenetic regulation of subgenome dominance following whole genome triplication in *Brassica rapa*. New Phytologist 211:288–299.

Cheng, H., Concepcion, G.T., Feng, X., Zhang, H., and Li, H. (2021). Haplotype-resolved de novo assembly with phased assembly graphs with hifiasm. Nature Methods 18:170–175.

Concia, L., Veluchamy, A., Ramirez-Prado, J.S., Martin-Ramirez, A., Huang, Y., Perez, M., Domenichini, S., Rodriguez Granados, N.Y., Kim, S., and Blein, T. (2020). Wheat chromatin architecture is organized in genome territories and transcription factories. Genome Biology 21:1–20.

Dobin, A., Davis, C.A., Schlesinger, F., Drenkow, J., Zaleski, C., Jha, S., Batut, P., Chaisson, M., and Gingeras, T.R. (2013). STAR: Ultrafast universal RNA-seq aligner. Bioinformatics 29:15–21. 10.1093/bioinformatics/bts635.

Dudchenko, O., Batra, S.S., Omer, A.D., Nyquist, S.K., Hoeger, M., Durand, N.C., Shamim, M.S., Machol, I., Lander, E.S., and Aiden, A.P. (2017). De novo assembly of the *Aedes aegypti* genome using Hi-C yields chromosome-length scaffolds. Science 356:92–95.

Edger, P.P., Poorten, T.J., VanBuren, R., Hardigan, M.A., Colle, M., McKain, M.R., Smith, R.D., Teresi, S.J., Nelson, A.D., and Wai, C.M. (2019). Origin and evolution of the octoploid strawberry genome. Nature Genetics 51:541–547.

Ellinghaus, D., Kurtz, S., and Willhoeft, U. (2008). LTRharvest, an efficient and flexible software for de novo detection of LTR retrotransposons. BMC Bioinformatics 9:18.

Emms, D.M., and Kelly, S. (2019). OrthoFinder: phylogenetic orthology inference for comparative genomics. Genome biology 20:1–14.

Gel, B., and Serra, E. (2017). karyoploteR: an R/Bioconductor package to plot customizable genomes displaying arbitrary data. Bioinformatics 33:3088–3090.

Ghamkhar, K., Croser, J., Aryamanesh, N., Campbell, M., Kon’kova, N., and Francis, C. (2010). Camelina (*Camelina sativa* (L.) Crantz) as an alternative oilseed: molecular and ecogeographic analyses. Genome 53:558–567.

Haas, B.J., Delcher, A.L., Wortman, J.R., and Salzberg, S.L. (2004). DAGchainer: a tool for mining segmental genome duplications and synteny. Bioinformatics 20:3643–3646.

He, X., Dias Lopes, C., Pereyra-Bistrain, L.I., Huang, Y., An, J., Chaouche, R.B., Zalzalé, H., Wang, Q., Ma, X., and Antunez-Sanchez, J. (2024). Genetic–epigenetic interplay in the determination of plant 3D genome organization. Nucleic Acids Research 52:10220–10234.

Higgins, E.E., Howell, E.C., Armstrong, S.J., and Parkin, I.A. (2020). A major quantitative trait locus on chromosome A9, *BnaPh1*, controls homoeologous recombination in *Brassica napus*. New Phytologist.

Hohmann, N., Wolf, E.M., Lysak, M.A., and Koch, M.A. (2015). A time-calibrated road map of Brassicaceae species radiation and evolutionary history. The Plant Cell 27:2770–2784.

Hollister, J.D., and Gaut, B.S. (2009). Epigenetic silencing of transposable elements: a trade-off between reduced transposition and deleterious effects on neighboring gene expression. Genome Research 19:1419–1428.

Holst, F., Bolger, A., Günther, C., Maß, J., Triesch, S., Kindel, F., Kiel, N., Saadat, N., Ebenhöh, O., and Usadel, B. (2023). Helixer–de novo prediction of primary eukaryotic gene models combining deep learning and a hidden Markov model. BioRxiv:2023.2002.2006.527280.

Hu, G., Grover, C.E., Vera, D.L., Lung, P.-Y., Girimurugan, S.B., Miller, E.R., Conover, J.L., Ou, S., Xiong, X., and Zhu, D. (2024). Evolutionary dynamics of chromatin structure and duplicate gene expression in diploid and allopolyploid cotton. Molecular Biology and Evolution 41:msae095.

Jin, J.-J., Yu, W.-B., Yang, J.-B., Song, Y., DePamphilis, C.W., Yi, T.-S., and Li, D.-Z. (2020). GetOrganelle: a fast and versatile toolkit for accurate de novo assembly of organelle genomes. Genome biology 21:1–31.

Kagale, S., Nixon, J., Khedikar, Y., Pasha, A., Provart, N.J., Clarke, W.E., Bollina, V., Robinson, S.J., Coutu, C., Hegedus, D.D., et al. (2016). The developmental transcriptome atlas of the biofuel crop *Camelina sativa*. Plant Journal 88:879–894. 10.1111/tpj.13302.

Kagale, S., Koh, C., Nixon, J., Bollina, V., Clarke, W.E., Tuteja, R., Spillane, C., Robinson, S.J., Links, M.G., Clarke, C., et al. (2014). The emerging biofuel crop *Camelina sativa* retains a highly undifferentiated hexaploid genome structure. Nature Communications 5:1–11. 10.1038/ncomms4706.

Katoh, K., and Standley, D.M. (2013). MAFFT multiple sequence alignment software version 7: improvements in performance and usability. Molecular biology and evolution 30:772–780.

Kaul, A., Bhattacharyya, S., and Ay, F. (2020). Identifying statistically significant chromatin contacts from Hi-C data with FitHiC2. Nature protocols 15:991–1012.

Kitamoto, N., Yui, S., Nishikawa, K., Takahata, Y., and Yokoi, S. (2014). A naturally occurring long insertion in the first intron in the *Brassica rapa FLC2* gene causes delayed bolting. Euphytica 196:213–223.

Krzywinski, M., Schein, J., Birol, I., Connors, J., Gascoyne, R., Horsman, D., Jones, S.J., and Marra, M.A. (2009). Circos: an information aesthetic for comparative genomics. Genome Research 19:1639–1645.

Kumar, S., Stecher, G., Li, M., Knyaz, C., and Tamura, K. (2018). MEGA X: molecular evolutionary genetics analysis across computing platforms. Molecular biology and evolution 35:1547–1549.

Kumar, S., Suleski, M., Craig, J.M., Kasprowicz, A.E., Sanderford, M., Li, M., Stecher, G., and Hedges, S.B. (2022). TimeTree 5: an expanded resource for species divergence times. Molecular biology and evolution 39:msac174.

Langmead, B., and Salzberg, S.L. (2012). Fast gapped-read alignment with Bowtie 2. Nature Methods 9:357–359. 10.1038/nmeth.1923.

Li, B., and Dewey, C.N. (2011). RSEM: accurate transcript quantification from RNA-Seq data with or without a reference genome. BMC Bioinformatics 12:323. 10.1186/1471-2105-12-323.

Li, H., and Durbin, R. (2009). Fast and accurate short read alignment with Burrows-Wheeler transform. Bioinformatics (Oxford, England) 25:1754–1760. 10.1101/gr.129684.111.

Li, P., Quan, X., Jia, G., Xiao, J., Cloutier, S., and You, F.M. (2016). RGAugury: a pipeline for genome-wide prediction of resistance gene analogs (RGAs) in plants. BMC genomics 17:1–10.

Lim, K.B., Yang, T.J., Hwang, Y.J., Kim, J.S., Park, J.Y., Kwon, S.J., Kim, J., Choi, B.S., Lim, M.H., and Jin, M. (2007). Characterization of the centromere and peri-centromere retrotransposons in *Brassica rapa* and their distribution in related *Brassica* species. The Plant Journal 49:173–183.

Luo, Z., Brock, J., Dyer, J.M., Kutchan, T.M., Augustin, M., Schachtman, D.P., Ge, Y., Fahlgren, N., and Abdel-Haleem, H. (2019). Genetic diversity and population structure of a *Camelina sativa* spring panel. Frontiers in Plant Science 10:184.

Lyons, E., Pedersen, B., Kane, J., and Freeling, M. (2008). The value of nonmodel genomes and an example using SynMap within CoGe to dissect the hexaploidy that predates the rosids. Tropical Plant Biology 1:181–190.

Mandáková, T., Pouch, M., Brock, J.R., Al-Shehbaz, I.A., and Lysak, M.A. (2019). Origin and evolution of diploid and allopolyploid *Camelina* genomes were accompanied by chromosome shattering. The Plant Cell 31:2596–2612.

Mao, Y. (2019). GenoDup Pipeline: a tool to detect genome duplication using the dS-based method. PeerJ 7:e6303.

Marçais, G., and Kingsford, C. (2011). A fast, lock-free approach for efficient parallel counting of occurrences of k-mers. Bioinformatics 27:764–770.

Martin, S.L., Lujan Toro, B., James, T., Sauder, C.A., and Laforest, M. (2022). Insights from the genomes of 4 diploid *Camelina* spp. G3 12:jkac182.

Martin, S.L., Smith, T.W., James, T., Shalabi, F., Kron, P., and Sauder, C.A. (2017). An update to the Canadian range, abundance, and ploidy of *Camelina* spp.(Brassicaceae) east of the Rocky Mountains. Botany 95:405–417.

Martin, S.L., Lujan-Toro, B.E., Sauder, C.A., James, T., Ohadi, S., and Hall, L.M. (2019). Hybridization rate and hybrid fitness for *Camelina microcarpa* Andrz. ex DC (♀) and *Camelina sativa* (L.) Crantz(Brassicaceae) (♂). Evolutionary Applications 12:443–455. 10.1111/eva.12724.

McKenna, A., Hanna, M., Banks, E., Sivachenko, A., Cibulskis, K., Kernytsky, A., Garimella, K., Altshuler, D., Gabriel, S., Daly, M., et al. (2010). The Genome Analysis Toolkit: A MapReduce framework for analyzing next-generation DNA sequencing data. Genome Research 20:1297–1303. 10.1101/gr.107524.110.20.

Okonechnikov, K., Conesa, A., and García-Alcalde, F. (2016). Qualimap 2: advanced multi-sample quality control for high-throughput sequencing data. Bioinformatics 32:292–294.

Ou, S., and Jiang, N. (2018). LTR_retriever: a highly accurate and sensitive program for identification of long terminal repeat retrotransposons. Plant Physiology 176:1410–1422.

Ou, S., Chen, J., and Jiang, N. (2018). Assessing genome assembly quality using the LTR Assembly Index (LAI). Nucleic acids research 46:e126–e126.

Ou, S., Su, W., Liao, Y., Chougule, K., Agda, J.R., Hellinga, A.J., Lugo, C.S.B., Elliott, T.A., Ware, D., and Peterson, T. (2019). Benchmarking transposable element annotation methods for creation of a streamlined, comprehensive pipeline. Genome biology 20:1–18.

Peakall, R., and Smouse, P.E. (2012). GenALEx 6.5: Genetic analysis in Excel. Population genetic software for teaching and research-an update. Bioinformatics 28:2537–2539. 10.1093/bioinformatics/bts460.

Poland, J.A., Brown, P.J., Sorrells, M.E., and Jannink, J.-L. (2012). Development of high-density genetic maps for barley and wheat using a novel two-enzyme genotyping-by-sequencing approach. PloS One 7:e32253.

Pritchard, J.K., Stephens, M., and Donnelly, P. (2000). Inference of population structure using multilocus genotype data. Genetics 155:945–959. 10.1111/j.1471-8286.2007.01758.x.

Qiao, X., Li, Q., Yin, H., Qi, K., Li, L., Wang, R., Zhang, S., and Paterson, A.H. (2019). Gene duplication and evolution in recurring polyploidization–diploidization cycles in plants. Genome biology 20:1–23.

Quinlan, A.R., and Hall, I.M. (2010). BEDTools: a flexible suite of utilities for comparing genomic features. Bioinformatics 26:841–842.

Ramírez-González, R., Borrill, P., Lang, D., Harrington, S., Brinton, J., Venturini, L., Davey, M., Jacobs, J., Van Ex, F., and Pasha, A. (2018). The transcriptional landscape of polyploid wheat. Science 361:eaar6089.

Robinson, J.T., Turner, D., Durand, N.C., Thorvaldsdóttir, H., Mesirov, J.P., and Aiden, E.L. (2018). Juicebox. js provides a cloud-based visualization system for Hi-C data. Cell systems 6:256–258.e251.

Schranz, M.E., Lysak, M.A., and Mitchell-Olds, T. (2006). The ABC’s of comparative genomics in the Brassicaceae: building blocks of crucifer genomes. Trends in Plant Science 11:535–542.

Scrucca, L., Fop, M., Murphy, T.B., and Raftery, A.E. (2016). mclust 5: clustering, classification and density estimation using Gaussian finite mixture models. The R Journal 8:289.

Servant, N., Varoquaux, N., Lajoie, B.R., Viara, E., Chen, C.-J., Vert, J.-P., Heard, E., Dekker, J., and Barillot, E. (2015). HiC-Pro: an optimized and flexible pipeline for Hi-C data processing. Genome biology 16:1–11.

Shi, J., and Liang, C. (2019). Generic Repeat Finder: a high-sensitivity tool for genome-wide de novo repeat detection. Plant physiology 180:1803–1815.

Sievers, F., Wilm, A., Dineen, D., Gibson, T.J., Karplus, K., Li, W., Lopez, R., McWilliam, H., Remmert, M., and Söding, J. (2011). Fast, scalable generation of high-quality protein multiple sequence alignments using Clustal Omega. Molecular systems biology 7:539.

Simão, F.A., Waterhouse, R.M., Ioannidis, P., Kriventseva, E.V., and Zdobnov, E.M. (2015). BUSCO: assessing genome assembly and annotation completeness with single-copy orthologs. Bioinformatics 31:3210–3212.

Singh, R., Bollina, V., Higgins, E.E., Clarke, W.E., Eynck, C., Sidebottom, C., Gugel, R., Snowdon, R., and Parkin, I.A. (2015). Single-nucleotide polymorphism identification and genotyping in *Camelina sativa*. Molecular Breeding 35:35.

Stevens, T.J., Lando, D., Basu, S., Atkinson, L.P., Cao, Y., Lee, S.F., Leeb, M., Wohlfahrt, K.J., Boucher, W., and O’Shaughnessy-Kirwan, A. (2017). 3D structures of individual mammalian genomes studied by single-cell Hi-C. Nature 544:59–64.

Su, W., Gu, X., and Peterson, T. (2019). TIR-learner, a new ensemble method for TIR transposable element annotation, provides evidence for abundant new transposable elements in the maize genome. Molecular plant 12:447–460.

Sun, H., Wu, S., Zhang, G., Jiao, C., Guo, S., Ren, Y., Zhang, J., Zhang, H., Gong, G., and Jia, Z.J.M.p. (2017). Karyotype stability and unbiased fractionation in the paleo-allotetraploid Cucurbita genomes. 10:1293–1306.

Tamura, K., Stecher, G., and Kumar, S. (2021). MEGA11: molecular evolutionary genetics analysis version 11. Molecular biology and evolution 38:3022–3027.

Tang, H., Woodhouse, M.R., Cheng, F., Schnable, J.C., Pedersen, B.S., Conant, G., Wang, X., Freeling, M., and Pires, J.C. (2012). Altered patterns of fractionation and exon deletions in *Brassica rapa* support a two-step model of paleohexaploidy. Genetics 190:1563–1574.

Team, R.C. (2021). R: A language and environment for statistical computing. R Foundation for Statistical Computing, Vienna, Austria. URL https://www.R-project.org.

Tepfer, M., Hurel, A., Tellier, F., and Jenczewski, E. (2020). Evaluation of the progeny produced by interspecific hybridization between *Camelina sativa* and *C. microcarpa*. Annals of Botany 125:993–1002.

Tirnaz, S., Zandberg, J., Thomas, W.J., Marsh, J., Edwards, D., and Batley, J. (2022). Application of crop wild relatives in modern breeding: An overview of resources, experimental and computational methodologies. Frontiers in Plant Science 13:1008904.

Vurture, G.W., Sedlazeck, F.J., Nattestad, M., Underwood, C.J., Fang, H., Gurtowski, J., and Schatz, M.C. (2017). GenomeScope: fast reference-free genome profiling from short reads. Bioinformatics 33:2202–2204.

Wang, L., Jia, G., Jiang, X., Cao, S., Chen, Z.J., and Song, Q. (2021). Altered chromatin architecture and gene expression during polyploidization and domestication of soybean. The Plant Cell 33:1430–1446.

Wang, M., Wang, P., Lin, M., Ye, Z., Li, G., Tu, L., Shen, C., Li, J., Yang, Q., and Zhang, X. (2018). Evolutionary dynamics of 3D genome architecture following polyploidization in cotton. Nature Plants 4:90–97.

Wang, S., Wang, H., Jiang, F., Wang, A., Liu, H., Zhao, H., Yang, B., Xu, D., Zhang, Y., and Fan, W. (2022). EndHiC: assemble large contigs into chromosome-level scaffolds using the Hi-C links from contig ends. BMC bioinformatics 23:1–19.

Wang, T., van Dijk, A.D., Zhao, R., Bonnema, G., and Wang, X. (2024). Contribution of homoeologous exchange to domestication of polyploid Brassica. Genome Biology 25:231.

Wang, Y., Tang, H., DeBarry, J.D., Tan, X., Li, J., Wang, X., Lee, T.-h., Jin, H., Marler, B., and Guo, H. (2012). MCScanX: a toolkit for detection and evolutionary analysis of gene synteny and collinearity. Nucleic acids research 40:e49–e49.

Woodhouse, M.R., Cheng, F., Pires, J.C., Lisch, D., Freeling, M., and Wang, X. (2014). Origin, inheritance, and gene regulatory consequences of genome dominance in polyploids. Proceedings of the National Academy of Sciences 111:5283–5288.

Xiong, W., He, L., Lai, J., Dooner, H.K., and Du, C. (2014). HelitronScanner uncovers a large overlooked cache of Helitron transposons in many plant genomes. Proceedings of the National Academy of Sciences 111:10263–10268.

Xu, Z., and Wang, H. (2007). LTR_FINDER: an efficient tool for the prediction of full-length LTR retrotransposons. Nucleic Acids Research 35:W265–W268.

Yang, M.H., and Fu, Y.B. (2017). AveDissR: An R function for assessing genetic distinctness and genetic redundancy. Applications in Plant Sciences 5:1700018.

Yang, Z. (1997). PAML: a program package for phylogenetic analysis by maximum likelihood. Computer applications in the biosciences 13:555–556.

Yang, Z. (2007). PAML 4: phylogenetic analysis by maximum likelihood. Molecular biology and evolution 24:1586–1591.

Yin, S., Wan, M., Guo, C., Wang, B., Li, H., Li, G., Tian, Y., Ge, X., King, G.J., and Liu, K. (2020). Transposon insertions within alleles of BnaFLC. A10 and BnaFLC. A2 are associated with seasonal crop type in rapeseed. Journal of Experimental Botany 71:4729–4741.

Zhang, L., Liu, L., Li, H., He, J., Chao, H., Yan, S., Yin, Y., Zhao, W., and Li, M. (2024). 3D genome structural variations play important roles in regulating seed oil content of *Brassica napus*. Plant Communications 5.

Zhang, M., Zhang, Y., Scheuring, C.F., Wu, C.-C., Dong, J.J., and Zhang, H.-B. (2012). Preparation of megabase-sized DNA from a variety of organisms using the nuclei method for advanced genomics research. nature protocols 7:467–478.

Zhang, R.-G., Wang, Z.-X., Ou, S., and Li, G.-Y. (2019). TEsorter: lineage-level classification of transposable elements using conserved protein domains. bioRxiv:800177.

Zhao, M., Zhang, B., Lisch, D., and Ma, J.J.T.P.C. (2017). Patterns and consequences of subgenome differentiation provide insights into the nature of paleopolyploidy in plants. 29:2974–2994.

Zhou, C., Brown, M., Blaxter, M., Consortium, D.T.o.L.P., McCarthy, S.A., and Durbin, R. (2024). Oatk: a de novo assembly tool for complex plant organelle genomes. bioRxiv:2024.2010.2023.619857.

