## Supplemental Figures S1-S14 for "A new subgenome of the *Camelina* genus reveals genome dominance is controlled by chromosomal proximity"

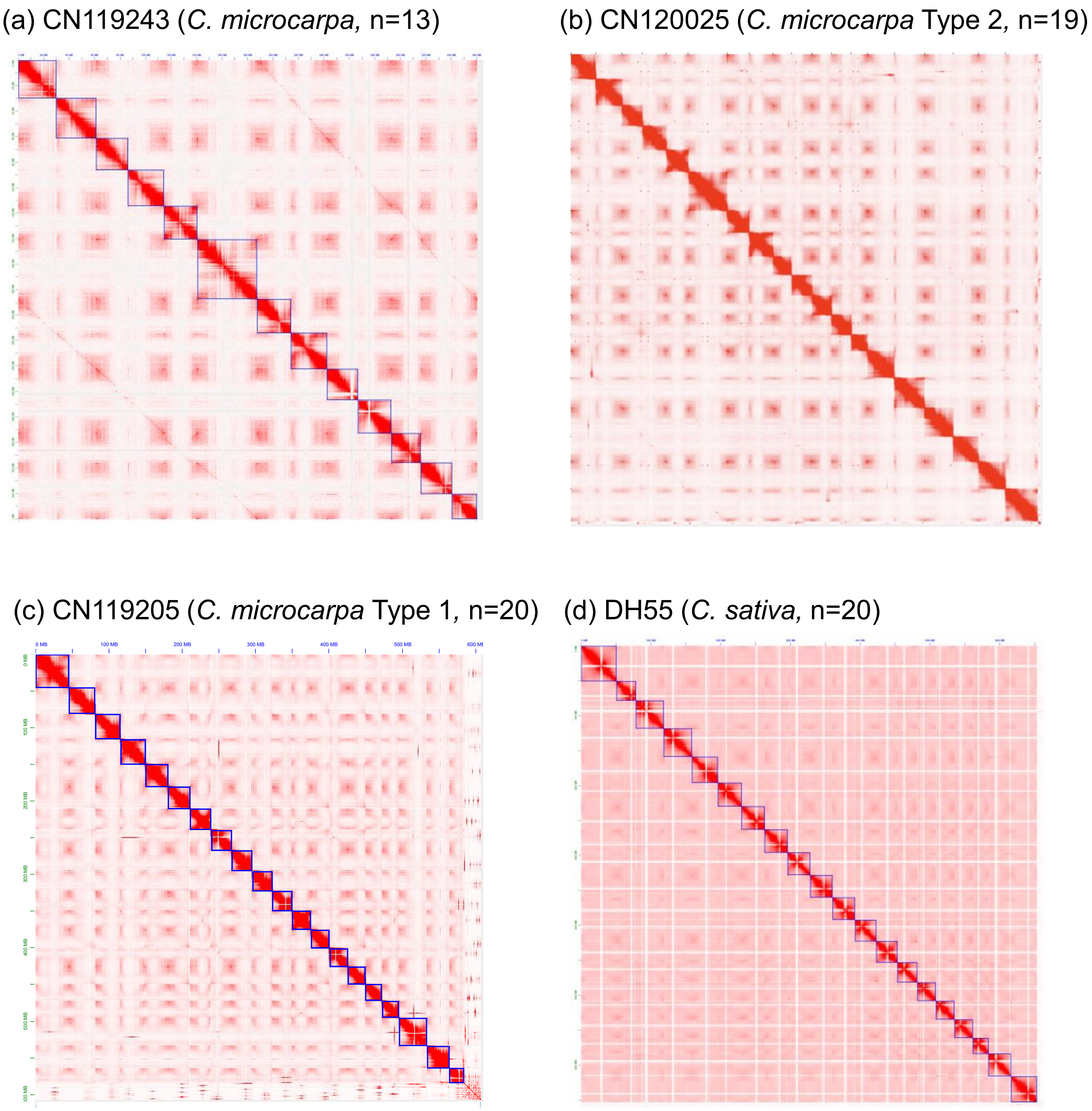


**Supplementary Figure 1**. Scaffolding of Camelina genomes using HiC technology

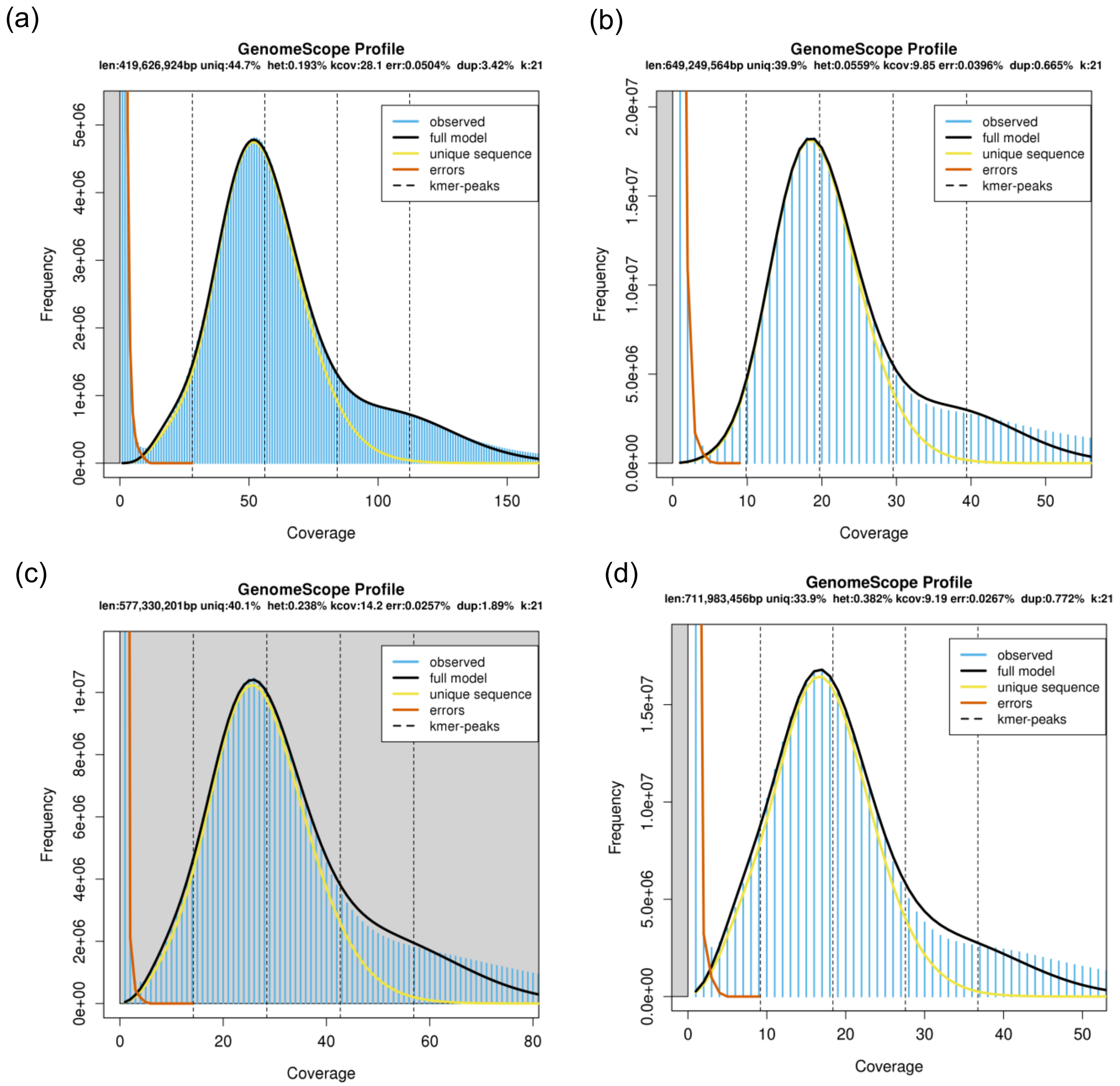


**Supplementary Figure 2**. Genome Size Estimation in Camelina species using kmer analysis *C. microcarpa* Tetraploid (a), *C. microcarpa* T1 (b), *C. microcarpa* T2 (c), and DH55 (d).

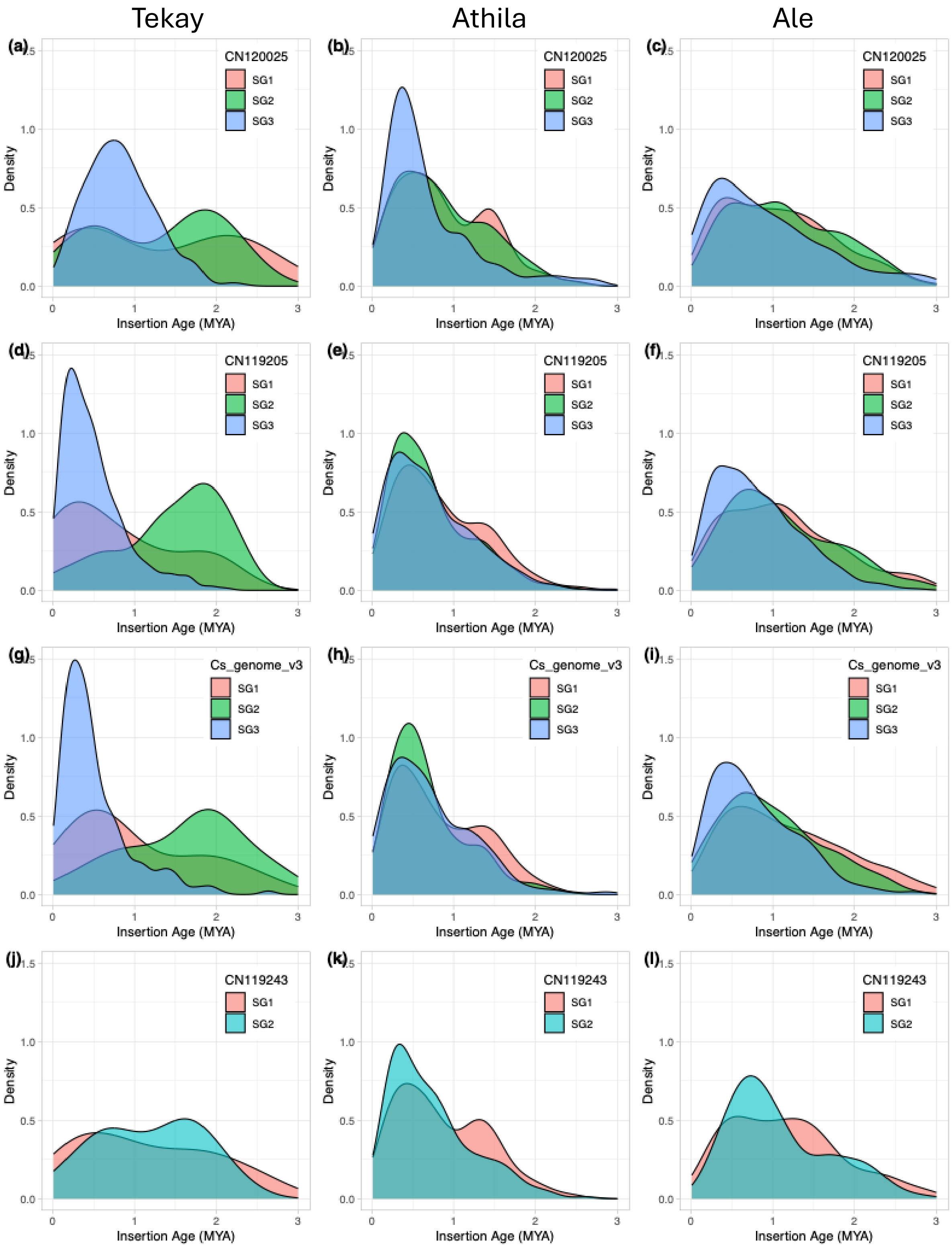


**Supplementary Figure 3**. Density of TE elements across hexaploid Camelina species. A. CmiT2 Tekay, B. Athila CmiT2, C. Ale CmiT2, D. Tekay CmiT1, E. Athila CmiT1, F. Ale CmiT1, G. Tekay CsaDH55, H. Athila CsaDH55, I Ale CsaDH55, J. Tekay Cmi4X, K. Athila Cmi4X, and L. Ale Cmi4X.

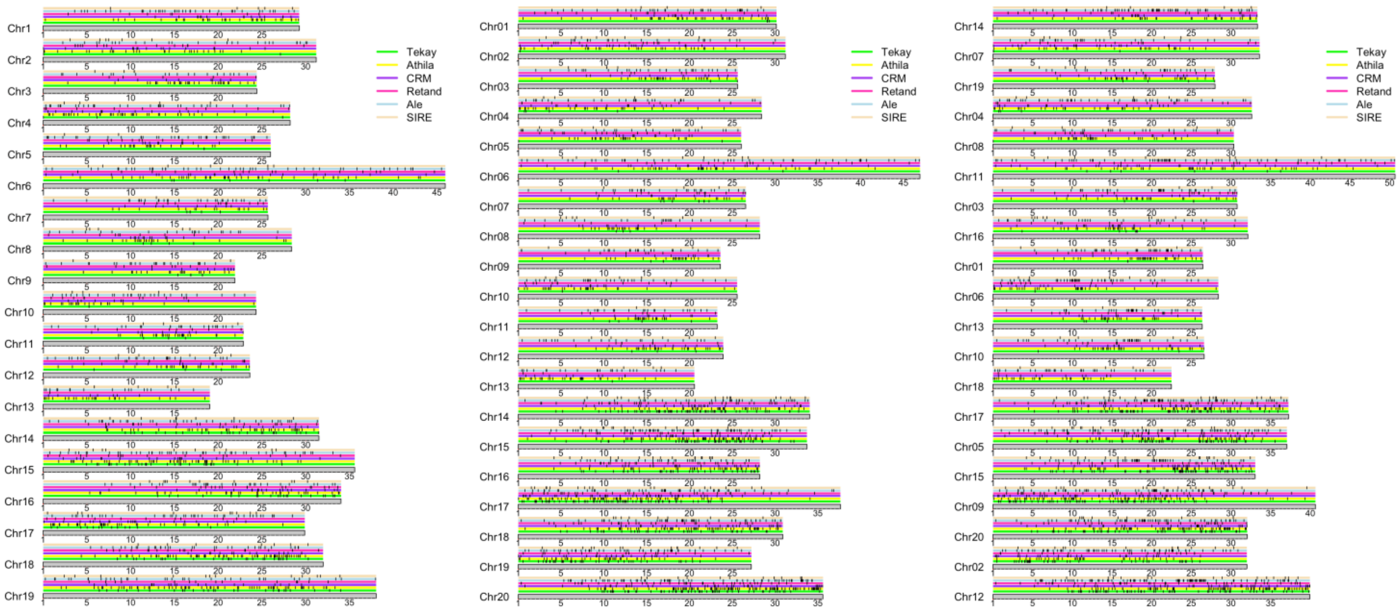


**Supplementary Figure 4**. Distribution of TEs across *Camelina* species (a) CmiT2; (b) CmiT1, and (c) CsaDH55. Different repeat types are represented in different tracks with different colours, each black dot represents the absolute position of a specific repeat.

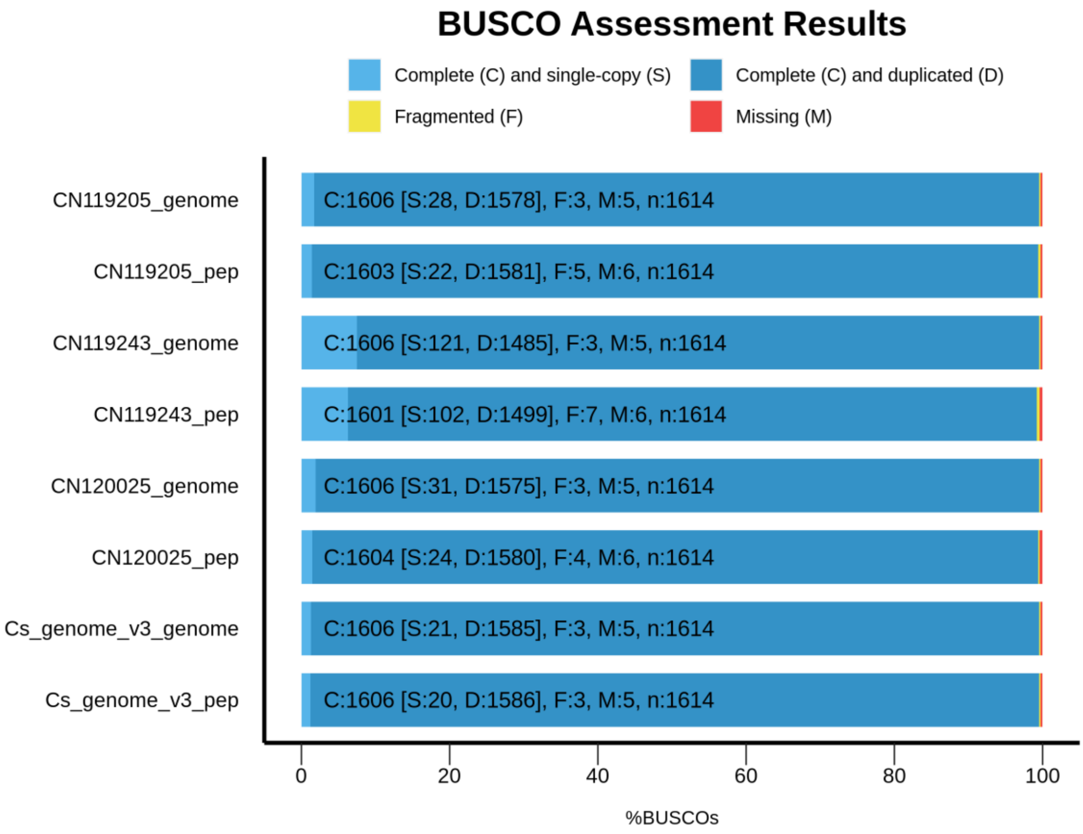


**Supplementary Figure 5**. BUSCO analysis of gene annotation for different Camelina species. CN119243 is Cmi4X, CN119205 is CmiT1, CN120025 is CmiT2, and Cs_genome_v3 is CsaDH55.

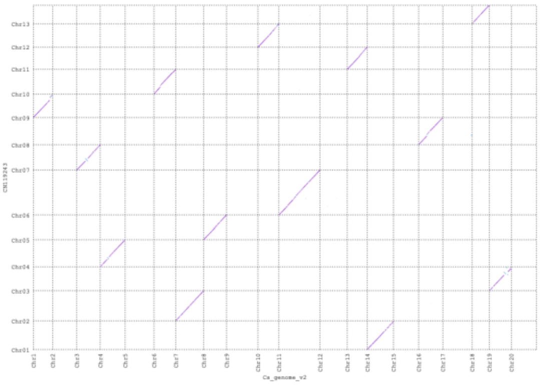

(a) DH55 - CN119243

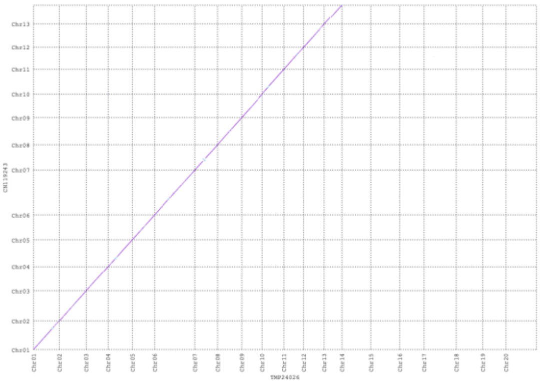

(b) CN119205 - CN119243

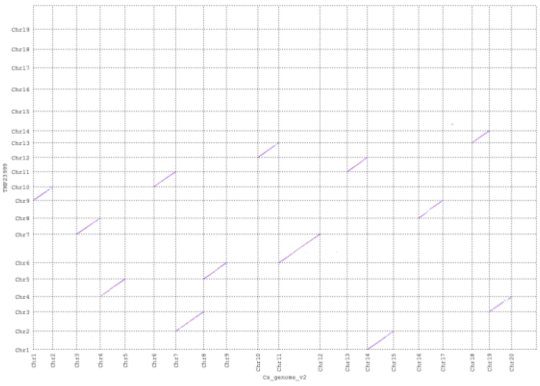

(c) DH55 - CN120025

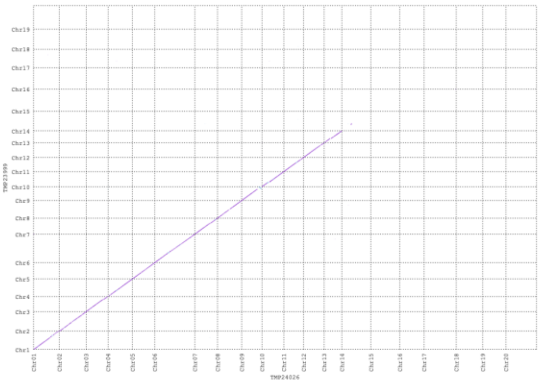

(d) CN119205 - CN120025

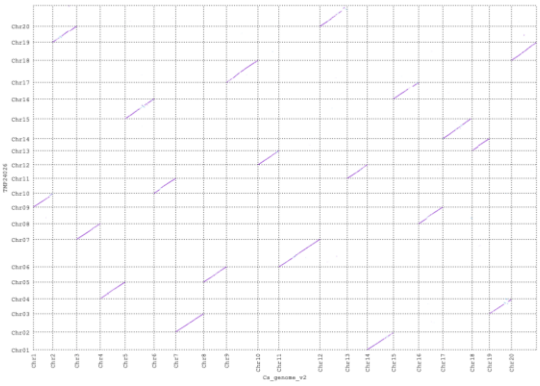

(e) DH55 - CN119205

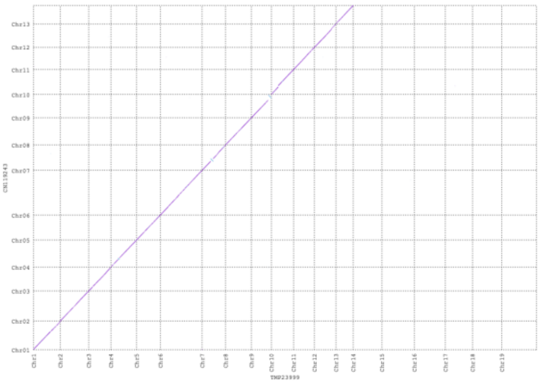

(f) CN120025 - CN119243

**Supplementary Figure 6**. Relationship among different species as explained by nucmer plots. CN119243 is Cmi4X, CN119205 is CmiT1, CN120025 is CmiT2, and DH55 is CsaDH55.

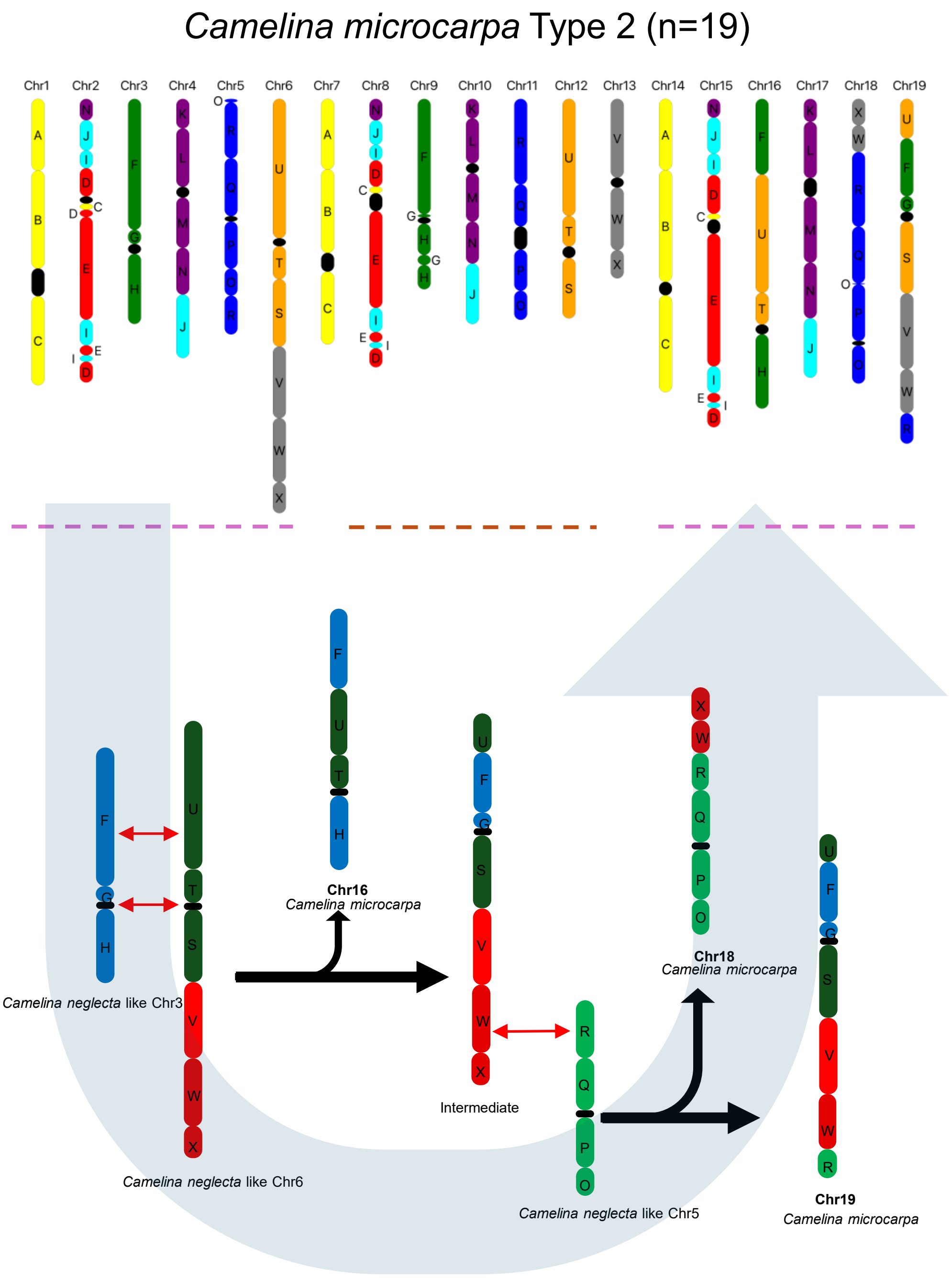


**Supplementary Figure 7**. Karyotype evolution in *C. microcarpa* Type 2

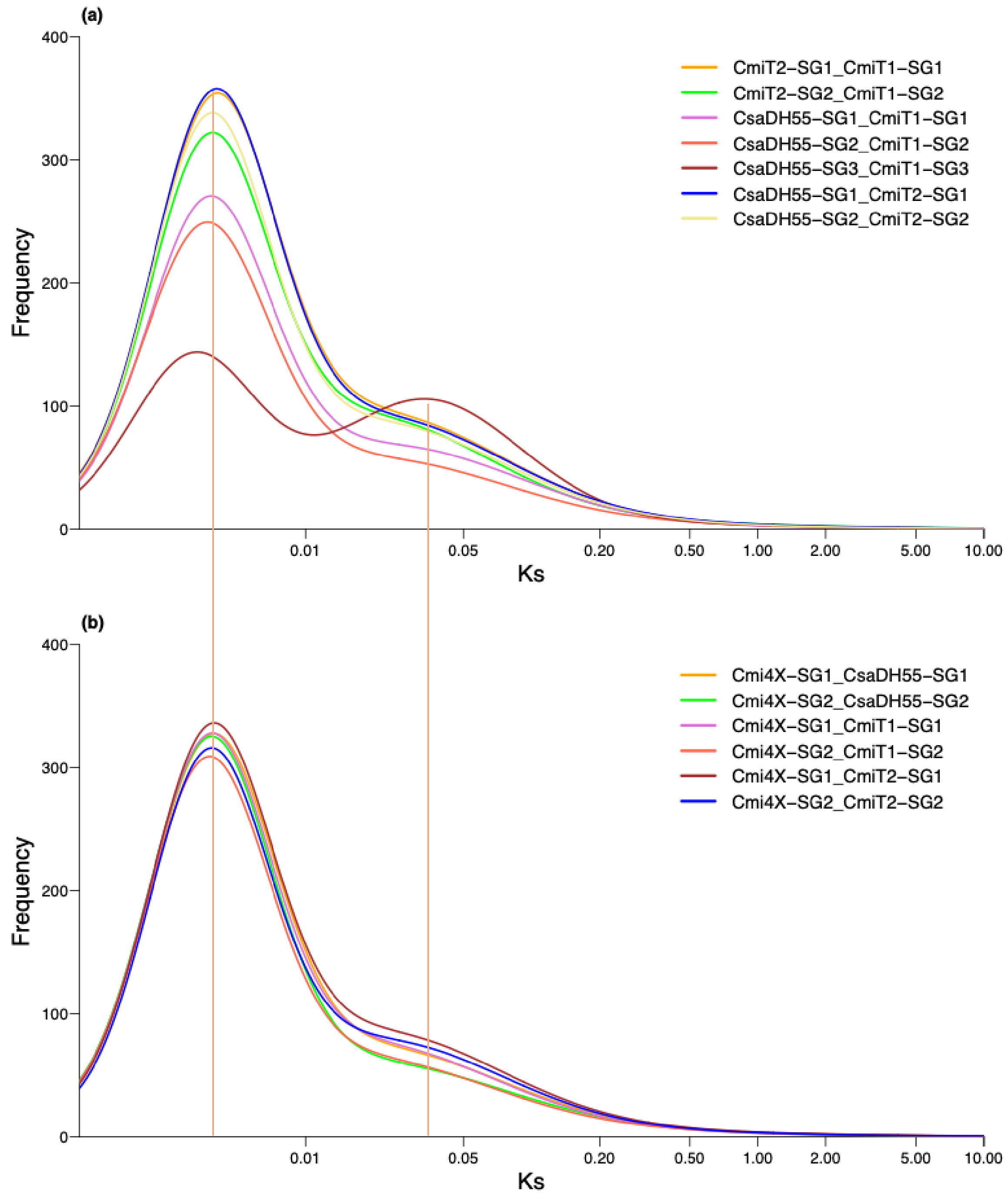


**Supplementary Figure 8**. Ks Analysis of subgenomes of hexaploid *Camelina* species (a) and Ks analysis of tetraploid *C. microcarpa* with subgenome of hexaploid *Camelina* species. CN119243 is Cmi4X, CN119205 is CmiT1, CN120025 is CmiT2, and CsaDH55 is *Camelina sativa*.

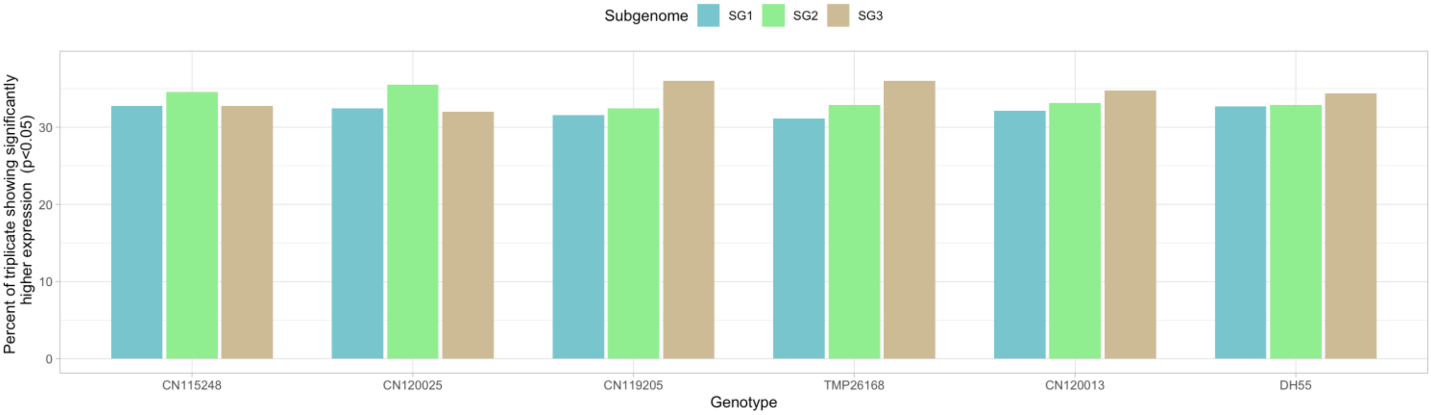


**Supplementary Figure 9**. Subgenome dominance analysis in *Camelina* species. The bar plot represents the percentage of genes showing significantly higher level of gene expression in each subgenome*. Camelina microcarpa* T2 is represented by CN115248 and CN120025; *C. microcarpa* T1 is represented by CN119205 and TMP26168; and *C. sativa* is represented by CN120013 and DH55.

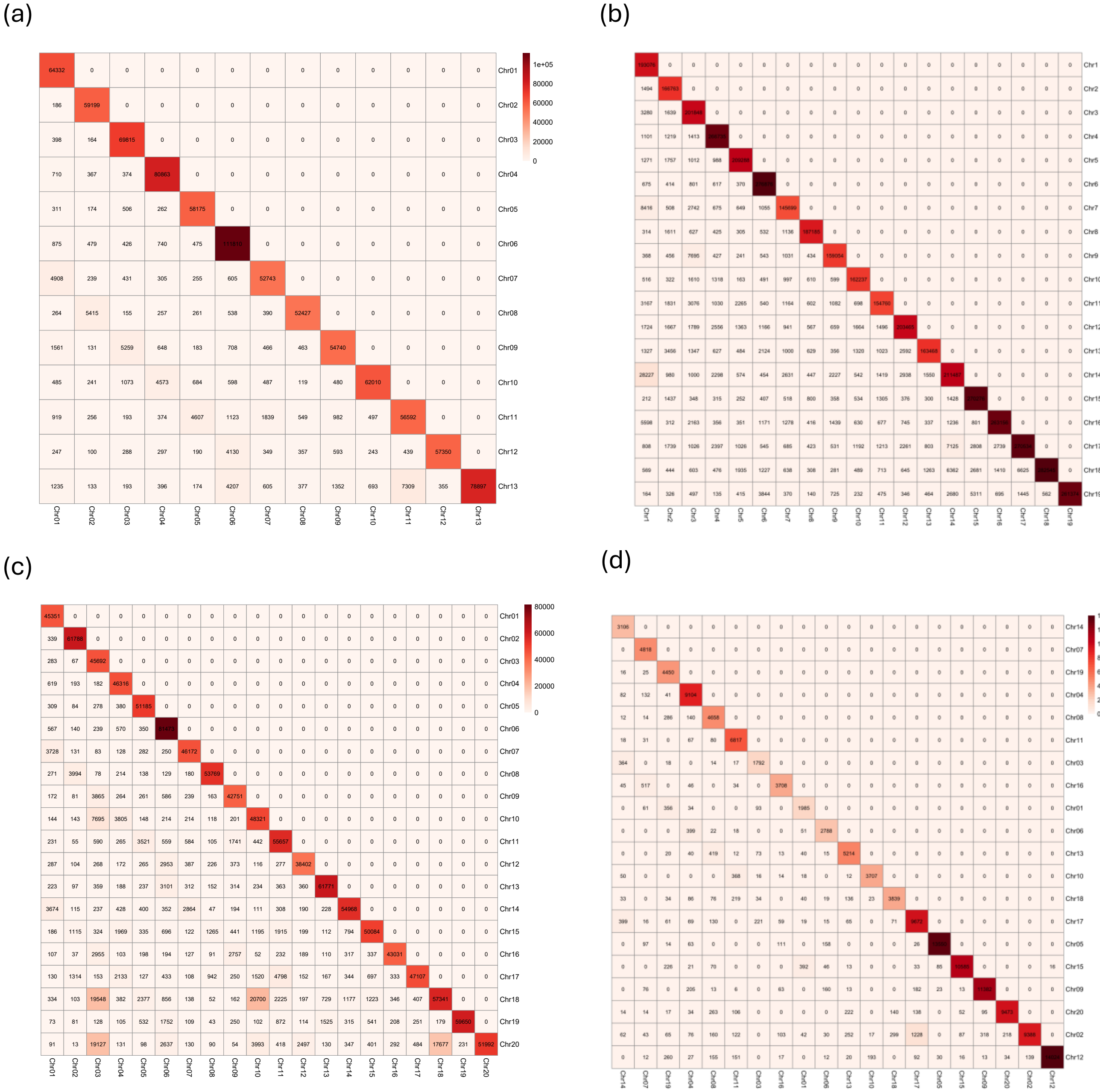


**Supplementary Figure 10**. Inter- and Intra-chromosomal Interaction as explained by chromatic capture confirmation methods in (a) CN119243, (b) CN120025, (c) CN119205, and (d) DH55 accession. CN119243 is Cmi4X, CN119205 is CmiT1, CN120025 is CmiT2, and DH55 is CsaDH55.

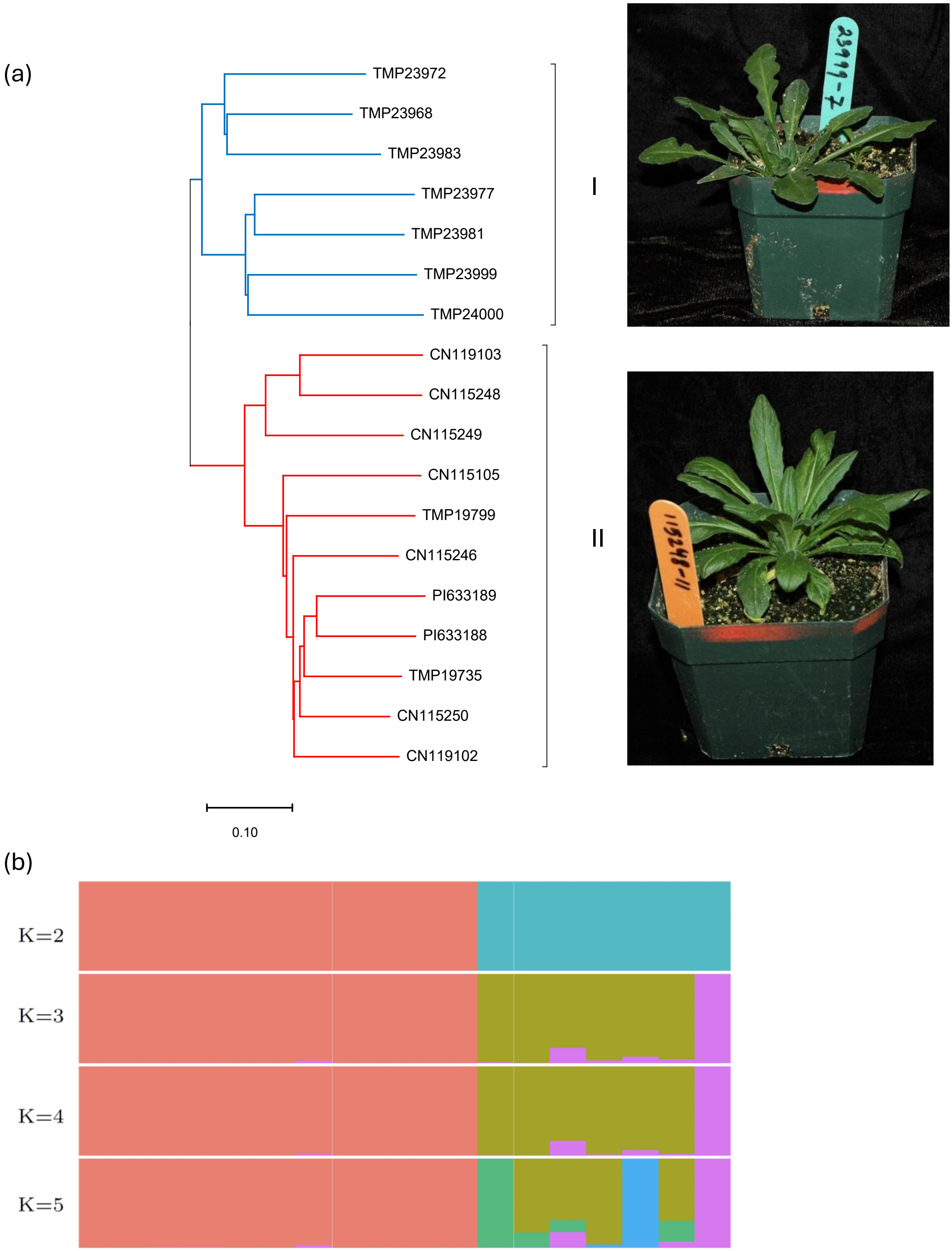


**Supplementary Figure 11**. Genetic relationship among 18 genotypes from *Camelina microcarpa*. NJ based phylogenetic tree where colour of branches represent genotypes in respective sub-populations (a), and population structure analysis with assignment probabilities (b).

(a) $F_2$ lines- 83-8

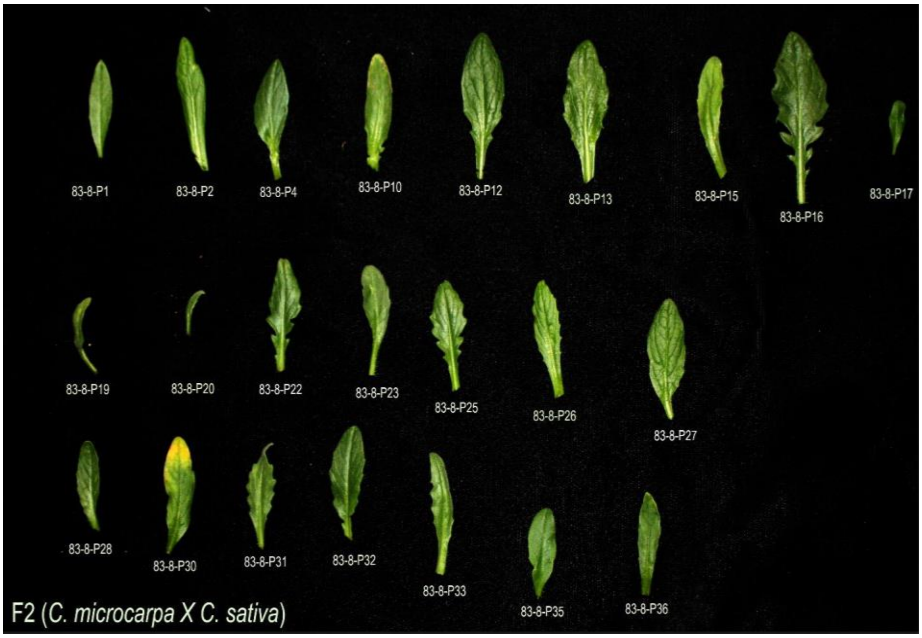


(b) $F_2$ lines-82-6

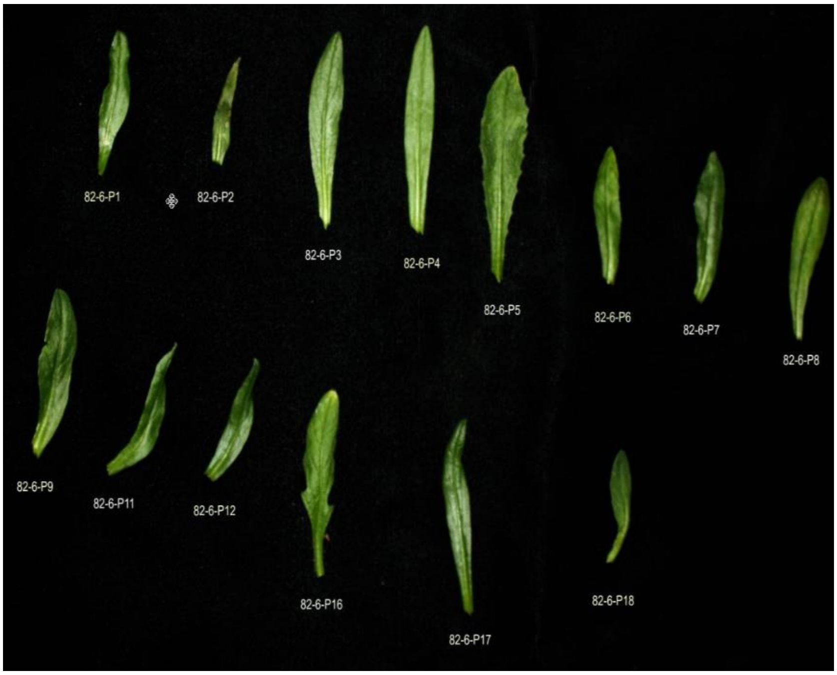


**Supplementary Figure 12**. Morphological variation in $F_2$ lines developed from interspecific hybridization between *C. microcarpa* Type 2 and *C. sativa.* The same parental lines were used to develop segregating lines; however, two different $F_1$ (a) 83-8 and (b) 82-6 were used to generated $F_2$ lines.

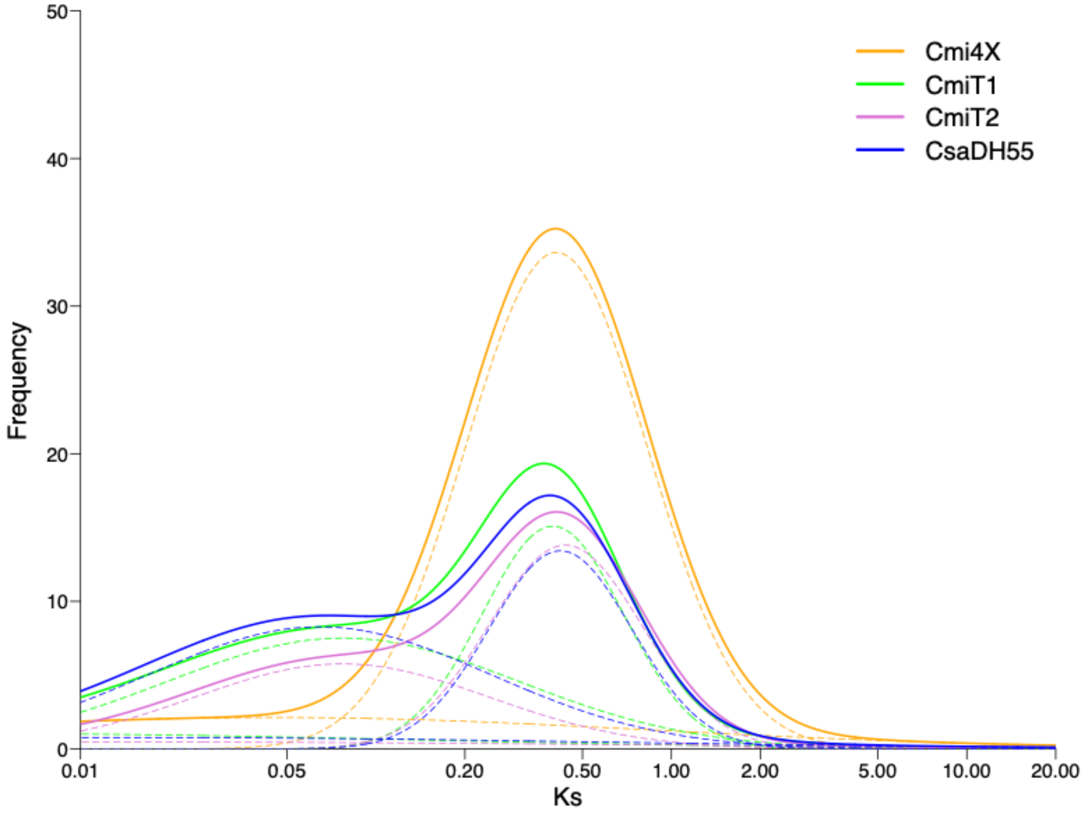


**Supplementary Figure 13**. Ks analysis with tandem duplicate genes in *Camelina* species. Dotted lines indicates different components to fit the gaussian model for the respective Ks values.

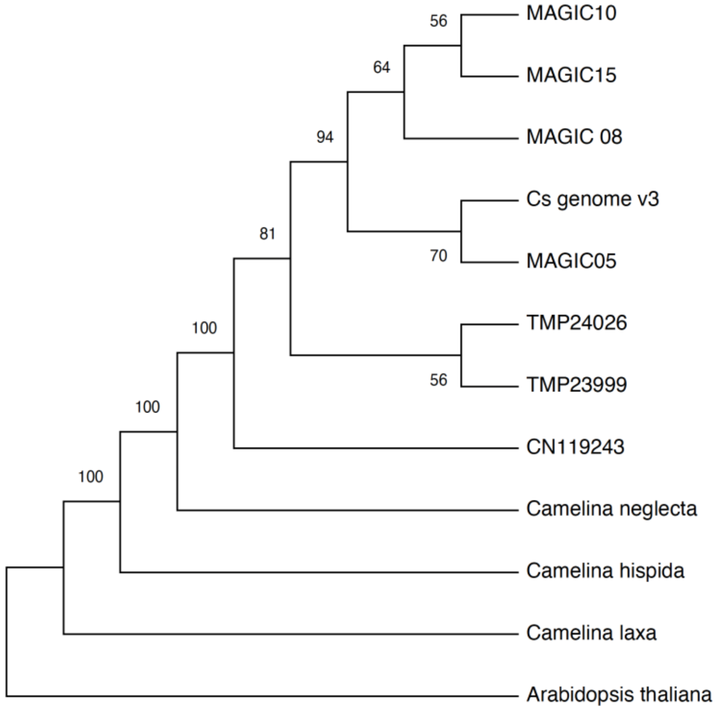


**Supplementary Figure 14**. Phylogenetic relationship among *Camelina* species based on whole chloroplast assembly. The *Camelina neglecta*, *C. laxa*, *C. hispida* were adopted from (Martin et al., 2022); CN119243 (*C. microcarpa* tetraploid (n=13), TMP24026 (*C. microcarpa* T1), TMP23999 (*C. microcarpa* T2), Cs genome v3 (DH55) were generated in this study, and MAGIC05, MAGIC08, MAGIC10 and MAGIC15 were unpublished genomes of *Camelina sativa*.
